# Parameter-wise predictions and sensitivity analysis for random walk models in the life sciences

**DOI:** 10.1101/2025.07.12.664534

**Authors:** Yihan Liu, David J. Warne, Matthew J. Simpson

## Abstract

Sensitivity analysis characterises input–output relationships for mathematical models, and has been widely applied to deterministic models across many applications in the life sciences. In contrast, sensitivity analysis for stochastic models has received less attention, with most previous work focusing on well-mixed, non-spatial problems. For explicit spatio-temporal stochastic models, such as random walk models (RWMs), sensitivity analysis has received far less attention. Here we present a new type of sensitivity analysis, called *parameter-wise prediction*, for two types of biologically-motivated and computationally expensive RWMs. To overcome the limitations of directly analysing stochastic simulations, we employ continuum-limit partial differential equation (PDE) descriptions as surrogate models, and we link these efficient surrogate descriptions to the RWMs using a range of biophysically-motivated *measurement error models*. Our approach is likelihood-based, which means that we also consider likelihood-based parameter estimation and identifiability analysis along with parameter sensitivity. The new approach is presented for two important classes of lattice-based RWM including a classical model where crowding effects are neglected, and an exclusion process model that explicitly incorporates crowding. Our workflow illustrates how different process models can be combined with different measurement error models to reveal how each parameter impacts the outcome of the expensive stochastic simulation. Open-access software to replicate all results is available on GitHub.

## 1 Introduction

Sensitivity analysis aims to examine how the output of a mathematical model relates to, or is influenced by the inputs. Model inputs can include model parameter values and initial conditions [1], while outputs typically refer to the solution of the mathematical model, the result of a single realisation of a stochastic model, or some average result obtained from an ensemble of stochastic simulations [2]. Many studies focus on deterministic models as they are widely used and easily interpretable in the sense that the model output is free from randomness and uncertainty [3–7]. Sensitivity analysis for stochastic models has been explored to a lesser extent, primarily for stochastic reaction networks (e.g. chemical reaction networks, birth–death models, gene expression networks) [8–12], where individuals within the population are assumed to be well-mixed and model outputs (e.g. the density of chemical species) are translationally invariant.

Lattice-based random walk models (RWMs), that describe the motion of individual agents on a lattice, are frequently used to interpret cell biology experiments involving cell migration and proliferation [13–19]. Here we focus on two types of RWMs, each motivated by a different kind of cell biology experiment. The first RWM is inspired by scratch assay data [20], using Fluorescence Ubiquitin Cell Cycle Indicator (FUCCI) labelling [21], illustrated in Figure 1(a), where cells in different phases of the cell cycle fluoresce red or green. A schematic of the cell cycle is illustrated in Figure 1(b). These experiments provide information about how cell position relates to cell cycle status [21], and are often performed under low-density conditions where cells are free to progress through the cell cycle without any contact inhibition [20, 22]. The second RWM is motivated by a circular barrier assay, illustrated in Figure 1(c), which involves growing high-density, uniform monolayers of cells within a circular barrier to mimic a high cell density *in vivo* environment [23, 24]. The circular barrier assay involves placing cells within the barrier and allowing time for cells to attach to the cell culture plate. The barrier is then lifted, non-viable cells are washed away, and the subsequent longer-term radial expansion of the population is imaged. Schematics of the circular barrier assay are illustrated in Figure 1(f)-(g).

**Figure 1.**
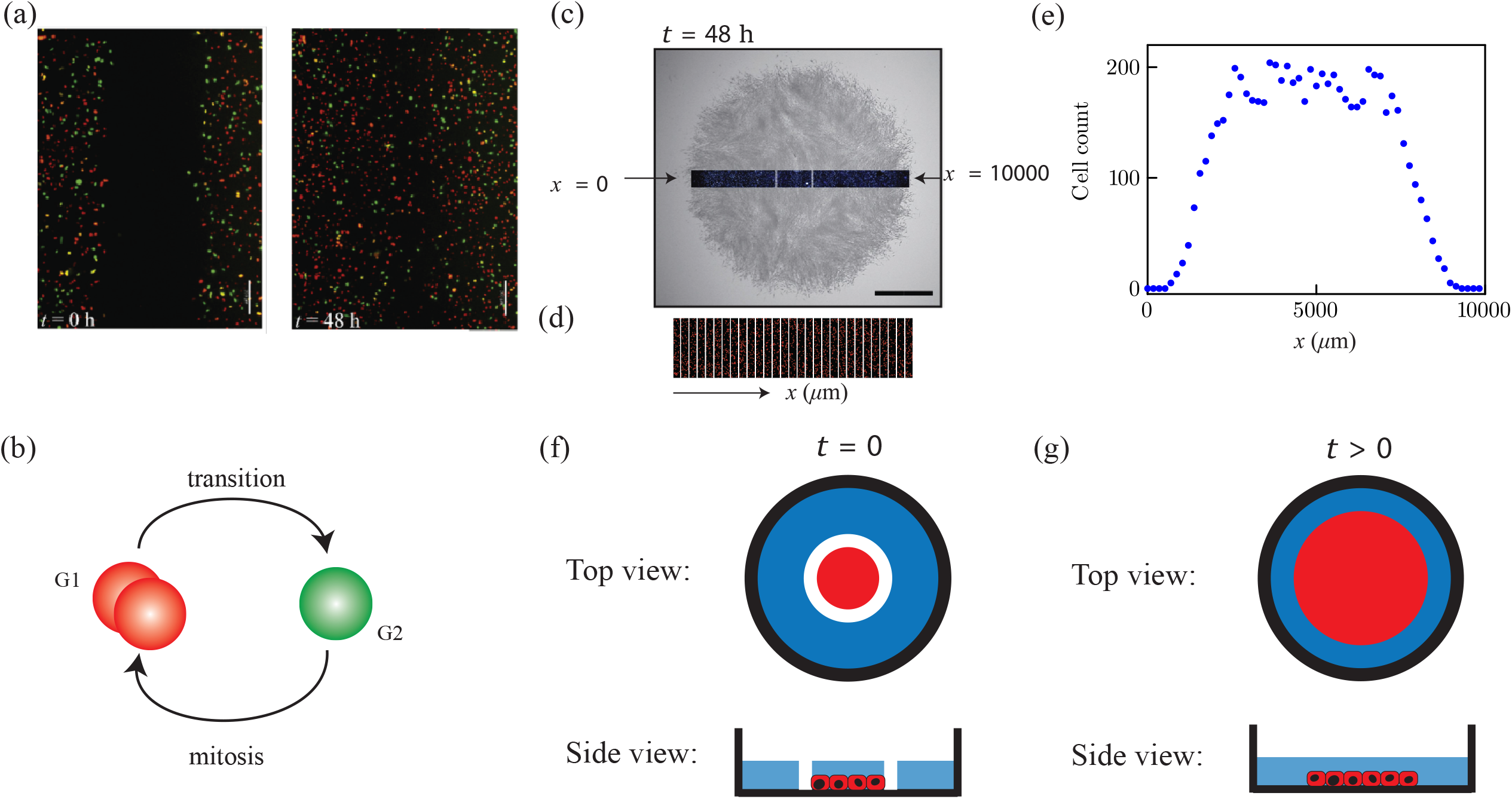
Experimental motivation. (a) Images of a scratch assay using melanoma cells with FUCCI at 0 h and 48 h. Scale bar corresponds to 200 µm [20]. (b) Cell cycle schematic indicating FUCCI fluorescence. (c) Barrier assay at 48 h with 2 mm scale bar [23]. (d) Transect through the centre of the spreading population in (c). (e) Count data collected from (d). (f)–(g) Schematic of the circular barrier assay. Black ring (open rectangle) represents the well, the blue region denotes the cell culture medium, the red regions indicate the expanding cell monolayer, and the white ring (bars) in (e) indicate the barrier. Images are reused with permission.

Parameter estimation and model predictions are important if we wish to gain mechanistic insight or to make predictions about future experimental scenarios. Estimation and prediction can be made using likelihood-based [13, 25–28] or Bayesian methods [17, 18, 29– 33]. For RWMs in the theoretical biology literature, it has become customary to focus on likelihood-free Bayesian approaches [17, 18, 29, 30, 33], but more recently we have focused on computationally efficient likelihood-based approaches that we will follow here [13].

In this work, we follow a likelihood-based approach by using maximum likelihood estimation (MLE) to estimate best-fit model parameters for a range of RWMs. To ensure these estimates are biologically meaningful and interpretable, we consider parameter identifiability analysis to determine whether the data contain sufficient information to yield precise estimates [3, 4, 13, 34–36]. In particular, we use the *profile likelihood* to assess the practical identifiability [35, 37–40] by partitioning the parameter vector *θ* into two parts, *θ* = (*ψ, ω*)^*⊤*^, where *ψ* represents the *interest* parameter(s) and *ω* represents the *nuisance* parameter(s) [37]. This targeting property of the profile likelihood allows us to assess parameter identifiability for complex models with many parameters by systematically isolating and analysing each parameter while optimising out the nuisance parameters. While predicting model outcomes across the full parameter space is routinely used to understand how uncertainties in parameter estimates propagate into uncertainties in model predictions [13, 41–43], this standard approach does not provide information about the role of individual parameters as in the context of sensitivity analysis that we focus on.

For RWMs in which agents migrate in physical space, sensitivity analysis has predominantly been conducted using methods originally designed for deterministic models, where the only source of uncertainty affecting the output is the variation in the input parameters and initial conditions [44]. Output uncertainties for RWMs arise from input variation (e.g. parameter values, initial condition) and from inherent stochasticity. To reduce the impact of inherent model stochasticity, model outputs are often averaged over many identically prepared realisations which can dramatically increase the computational cost of the analysis [45–48]. Of the few studies that perform sensitivity analyses for RWMs, most employ the partial rank correlation coefficient (PRCC) [45–48]. The PRCC quantifies the influence of each input parameter through a numerical index, called the *sensitivity index* [44, 49]. A positive (negative) sensitivity index indicates a positive (negative) correlation between the parameter and the model output. Sensitivity analysis using PRCC is typically combined with Latin hypercube sampling (LHS) [50]. To perform LHS-PRCC, a distribution is specified for each input parameter, and LHS is used to generate random parameter combinations that cover the parameter space. For each random parameter combination, the model is simulated to produce the model output(s). All input and outputs are rank-transformed. To isolate the impact of a target input, two linear regressions are performed: (i) the target input is regressed on the remaining inputs, and (ii) the output is regressed on all inputs except the target input. The sensitivity index is calculated as the correlation between the residuals from the two regressions. While LHS-PRCC can isolate the effect of parameters one at a time, randomly drawn parameter samples can lead to working with insufficient or implausible parameter combinations [44] which means that LHS-PRCC can be computationally inefficient. Therefore, addressing the limitations of working with random parameter samples by developing computationally efficient approaches for sensitivity analysis remains a significant challenge in the field of RWMs.

One approach to reduce the computational cost of sensitivity analysis is to use efficient surrogate models to approximate the output of a RWM [51–54]. This can include working with continuum-limit descriptions [13, 41, 51, 55, 56]. In addition to improving computational efficiency via working with surrogates, we also avoid working with randomly chosen sample parameters as in LHS-PRCC. To this end, Simpson and Maclaren [3, 57] proposed an efficient method for sensitivity analysis based on the targeting property of the profile likelihood. Parameter-wise prediction intervals provide both qualitative and quantitative insight into how uncertainty in individual parameters, or combinations of parameters, influences variability in model predictions. Importantly, the parameter combinations used to construct parameter-wise prediction intervals are not random combinations. Instead, parameter combinations are based on optimising a log-likelihood function which ensures that we focus on relevant parameter combinations that lead to model solutions that are close to the observed data. The parameter-wise prediction approach has been applied to various continuum models, including both identifiable [3, 4, 57] and non-identifiable models [34], but it has not yet been employed in the context of more complicated and computationally expensive stochastic models, such as RWMs. In this work, we conduct the first sensitivity analysis for RWMs using parameter-wise prediction.

## 2 Methods

### 2.1 Motivating experiments

We use two types of lattice-based RWMs motivated by two different kinds of cell biology experiments presented schematically in Figure 1. The first experiment is a scratch assay performed under low cell density conditions, as illustrated in Figure 1(a) [20]. Scratch assays involve growing uniform monolayers of cells on tissue culture plates before part of the monolayer is scratched away. The resulting temporal recolonisation of the scratched region is imaged [58, 59]. The scratch assay of interest is conducted using FUCCI labelling where cells in G1 phase fluoresce red, and cells in G2 phase fluoresce green [20]. Cells in G1 phase progress through the cell cycle into G2 phase, and cells in G2 phase undergo mitosis, producing two daughter cells that are both in G1 phase, see Figure 1(b). These experiments are performed under low density conditions to minimise the impact of contact inhibition so it is natural to model this experiment using a non-interacting RWM [19]. The RWM is non-interacting in the sense that multiple agents can reside on the same site, and agent movement is unaffected by the location of other agents. Since the experiments involve two subpopulations, each of which corresponds to cells in different phases of the cell cycle, the non-interacting RWM will also involve two subpopulations that represent cells in the two different phases of the cell cycle as identified in the experiments.

The second experiment is a circular barrier assay conducted under high cell density conditions, describing the spatial expansion of a circular monolayer of cells, as illustrated schematically in Figure 1(f)–(g). In this experiment, circular barriers are placed at the centre of each well in a 24-well tissue culture plate. Cells are seeded uniformly within circular barriers, and some time is allowed for cells to attach to the culture plate before the barriers are removed to initiate the experiment at *t* = 0. The initial cell monolayer is briefly washed to remove non-viable and unattached cells, and the subsequent outward radial expansion of the cell population is imaged at later times, as shown in Figure 1(c). As the circular barrier assays are conducted under high cell density conditions, it is important to account for crowding effects when modelling these experiments so we use an interacting RWM. The RWM is interacting in the sense that each lattice site can be occupied by, at most, a single agent, and agent movement is affected by the location of other agents on the lattice [19].

### 2.2 Mathematical models

Sections 2.2.1–2.2.2 present details of the non-interacting and interacting RWMs, respectively. Both models involve a 2D square lattice with lattice spacing Δ, and individual events are simulated using a random sequential update method [60]. In all simulations, each agent represents an individual biological cell. The lattice has dimensions *I × J*, consisting of *I* columns and *J* rows, with reflecting boundary conditions applied along all boundaries. Each lattice site is indexed by (*i, j*), and is associated with a unique Cartesian coordinate (*x*_*i*_, *y*_*j*_), where *x*_*i*_ = (*i* − 1)Δ for *i* = 1, 2, 3, …, *I*, and *y*_*j*_ = (*j* − 1)Δ for *j* = 1, 2, 3, …, *J*. All RWMs are discrete-time models, and the evolution of the models involves time steps of constant duration, *τ*. Time steps are indexed by *k* so that *t*_*k*_ = *kτ* for *k* = 1, 2, 3, …, *K*. To keep our simulation framework general, we work with dimensionless simulations by setting Δ = *τ* = 1. These simulations can be re-scaled using appropriate length and time scales to accommodate cells of different sizes, motility rates, and cell cycle progression rates [61].

#### 2.2.1 Non-interacting random walk model

Let *R*(*t*) and *G*(*t*) denote the total number of red and green agents that represent cells in G1 and G2 phase at time *t*, respectively, so that the total number of agents is *T* (*t*) = *R*(*t*)+*G*(*t*). We initialise the RWM by placing *R*(0) red agents, and *G*(0) green agents on the lattice. Since the RWM is non-interacting, each site can be occupied by multiple agents. To evolve the model from time *t* to time *t*+*τ*, we consider four types of stochastic events: (i) motility of red agents; (ii) motility of green agents; (iii) red agents progressing through the cell cycle and transitioning into green agents (red-to-green transition); and, (iv) green agents undergoing mitosis to produce two daughter agents that will be at the beginning of their cell cycle, belonging to the G1 subpopulation and accordingly coloured red.

To simulate agent motility, *R*(*t*) red agents and *G*(*t*) green agents are selected independently at random, one at a time, with replacement. Each selected agent is given an opportunity to move. Experimental evidence suggests motility of cells in G1 phase can be indistinguishable from cells in G2 phase [62]. We incorporate this into our model by allowing each selected agent to move with probability *P* ∈ [0, 1], regardless of their cell cycle status. The target site for a potential motility event is chosen as follows: the probability that a motile agent at site (*i, j*) steps to (*i, j ±*1) is 1*/*4, while the probability that it steps to (*i±*1, *j*) is (1*±ρ*)*/*4, where *ρ* ∈ [−1, 1] is a bias parameter that influences directional bias in the horizontal direction. Again, the bias parameter is taken to be identical for both subpopulations. Setting *ρ* = 0 corresponds to unbiased motility. The bias is considered only in the horizontal direction, as the macroscopic density of both subpopulations is independent of vertical location due to the geometry of the experiment in Figure 1(a). While the scratch assay in Figure 1(a) does not provide any obvious visual evidence of migration bias, it is often thought that bias mechanisms (e.g. chemotaxis) can play important roles under certain circumstances [63, 64]. For completeness, we include the bias term in our model so it is clear that our approach is applicable to both biased and unbiased cell migration. To simulate the red-to-green transitions, *R*(*t*) red agents are selected independently at random, one at a time, with replacement. Each selected red agent progresses through the cell cycle, transitioning into a green agent with probability *λ*_1_ ∈ [0, 1]. To simulate proliferation, *G*(*t*) green agents are selected independently at random, one at a time, with replacement. Each selected green agent undergoes mitosis to produce two red agents with probability *λ*_2_ ∈ [0, 1]. When proliferation takes place, both red daughter agents are placed on the same lattice site as the original parent green agent.

Although cells are free to move in any direction in the experiment in Figure 1(a), the macroscopic cell density is independent of the vertical position and varies with horizontal position. This geometric simplification is due to the fact that scratches are made within uniform monolayers. Experimental images within this setting can be quantified using count data obtained by superimposing a series of uniformly spaced columns across the image and counting the number of red and green cells separately in each column [41]. This approach summarises the experimental outcome as a series of count data as a function of the horizontal location of each column, as in Figure 1(c)–(e). For simplicity, it is common to report count data at a single time point at the end of the experiment, and we model this approach by using our models to simulate count data at a single time point only.

We will now explain how to generate count data from the non-interacting RWM so that it mimics the geometry and count data collected from the experiment in Figure 1(a). The experiment is designed to allow the population of cells to migrate inwards into an initially-vacant region, where the moving fronts eventually meet and the scratched region becomes occupied, as in Figure 1(a). In contrast, our RWM is designed such that agents tend to move outwards into empty space, resulting in a persistent spreading population with leading edges that progress outward without meeting. This outward migration design provides more information, particularly for simulations over longer time periods [65]. Despite these differences in movement direction, the important geometric features of our RWM match the experiment. All simulations are initialised such that the expected occupancy of lattice sites within each column is identical. Combined with reflecting boundary conditions, this ensures that the average density of any lattice site remains independent of vertical position throughout the simulation as in the experiment. The outcome of any RWM simulation provides the number of agents at each lattice site for different subpopulations as a function of time. We define 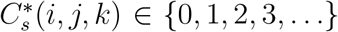 as the unbounded integer number of agents from the *s*th subpopulation at site (*i, j*) after *k* time steps, where *s* = 1, 2, …, *S*. For our application, we have *S* = 2. With this notation, the total number of agents from subpopulation *s* in the *i*th column after *k* time steps is

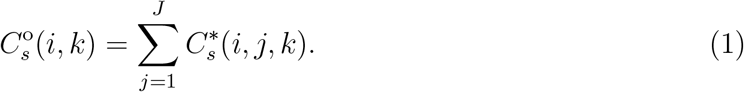

While the RWM provides a high-fidelity means of generating noisy data that mimic those collected from cell biology experiments, it can be computationally expensive to generate large numbers of forward simulations to obtain detailed information about the impact of different parameter choices. To address this limitation, we use a computationally efficient continuum-limit surrogate model that is given by

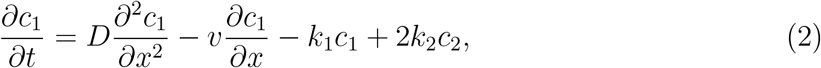

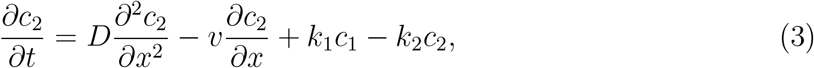

Where

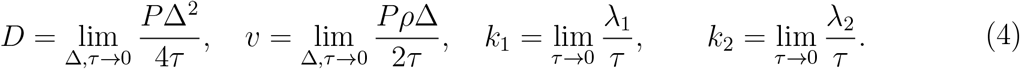

Here, *c*_1_(*x, t*) *≥* 0 and *c*_2_(*x, t*) *≥* 0 are the dimensionless densities of red and green agents at location *x* and time *t*, respectively. For the geometry and initial condition that we consider, Equations (2)–(3) have an exact solution for which the details are provided in Section 1 of the Supplementary Material.

An implicit assumption in deriving the surrogate continuum-limit model is that we are working with a sufficiently large lattice so that the discrete count data can be approximated by a smooth continuous function of position. Under this idealised scenario, it is possible to precisely relate the count data generated from the RWM to the solution of the corresponding continuum-limit PDE model in the following way,

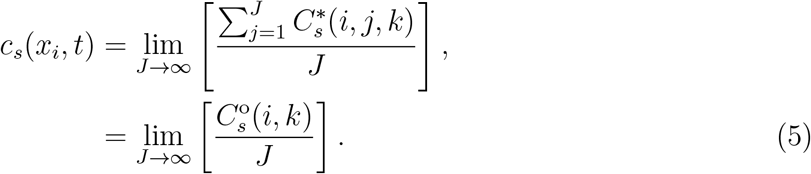

This idealised relationship is confirmed in Section 3 of the Supplementary Material. In practice however, we must deal with the case that *J* is finite, as experiments always involve a relatively small field of view, corresponding to small, finite *J*. These practical conditions mean that count data typically exhibits large fluctuations, such as the experimental count data in Figure 1(e). Consequently, the relationship between the observed count data, 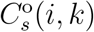, and the solution of the continuum-limit model, *c*_*s*_(*x, t*), is unclear. We will address this question by introducing physically-motivated *measurement error models* for different kinds of RWMs in Section 2.4 [4]. We note that measurement error models are also referred to as error models or observation models in the literature. In this work we use the term measurement error model.

#### 2.2.2 Interacting random walk model

Simulations of circular barrier assays are initialised by uniformly placing *T* (0) agents on lattice sites within a circular region of radius *r*_0_. Each site within this circular region has an occupancy probability *c*_0_ ∈ [0, 1] and since we are dealing with an interacting RWM each site can be occupied by, at most, a single agent. To evolve the stochastic algorithm from time *t* to time *t* + *τ*, we consider two types of stochastic events: (i) unbiased agent motility; and, (ii) agent proliferation. Motility events are simulated in a similar way to those in the non-interacting RWM, except that here we have a single population, *S* = 1, and we set *ρ* = 0, since our experimental images indicate the outward spreading population maintains a circular shape. To simulate proliferation, *T* (*t*) agents are selected independently at random, one at a time, with replacement, and each is given an opportunity to proliferate. When an agent is selected, it attempts to proliferate with probability *λ* ∈ [0, 1]. Upon proliferation, the agent deposits a daughter agent at one of its four nearest neighbour sites with equal probability. The key difference between the non-interacting and interacting RWM is that here we account for crowding effects by aborting potential motility or proliferation events that would place any agent on an occupied site. This means that the interacting RWM is closely related to an exclusion process [60].

The outcome of the circular barrier assay in Figure 1(c) is quantified by placing a narrow rectangular transect across the spreading monolayer [23]. Count data are collected by superimposing a series of uniformly spaced columns across the transect, as illustrated in Figure 1(d). We summarise the experimental data as a series of cell counts as a function of horizontal position, as shown in Figure 1(e). The interacting RWM can be used to generate count data that closely resembles the experimental data. To generate count data we place a rectangular transect across the centre of the lattice. The transect has dimensions *I ×H*, with the lower horizontal boundary at *j* = *j*_1_ and the upper horizontal boundary at *j* = *j*_1_ + *H*. We let *C*^*\**^(*i, j, k*) represent the occupancy of site (*i, j*) after *k* time steps. If site (*i, j*) is occupied we have *C*^*\**^(*i, j, k*) = 1, otherwise *C*^*\**^(*i, j, k*) = 0. The count data at the *i*th column after *k* time steps within the transect is

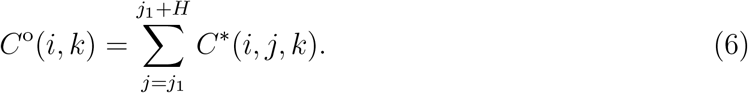

so that *C*°(*i, k*) ∈ {0, 1, 2, 3, …, *H*}. In summary, a key difference between the two types of column-based count data that we consider in this work is that count data from the non-interacting RWM are unbounded 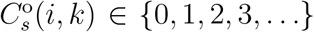, while count data from the interacting RWM are bounded *C*°(*i, k*) ∈ {0, 1, 2, 3, …, *H*}.

The continuum-limit description of the interacting RWM is a 2D generalisation of the well-known Fisher–Kolmogorov equation [61, 66], given by

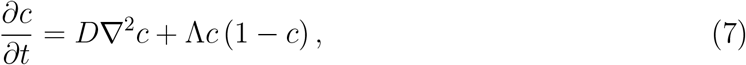

Where

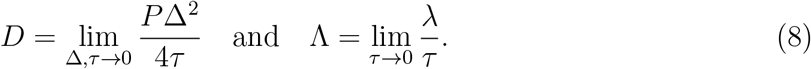

Since both the experiment and simulation maintain radial symmetry, we write Equation (7) as

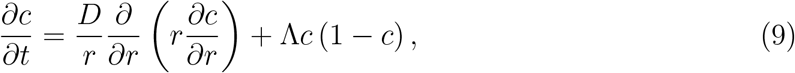

where *c*(*r, t*) *≥* 0 represents the dimensionless cell density at radius *r* and time *t*. Since Equation (9) cannot be solved analytically, we obtain numerical solutions as outlined in Section 2 of the Supplementary Material. For the simulations we consider, the computational time required to solve Equation (9) is approximately three orders of magnitude faster than simulating the RWM. Therefore, despite working with a numerical approach, the surrogate PDE model is very attractive for the purposes of estimation, identifiability and sensitivity analysis over working with the stochastic model. To link the solution of the continuum-limit model in Equation (9) with count data from the interacting RWM, we make the assumption that the agent density is constant within each rectangular discretisation along the transect. This will be a reasonable approximation provided that *H* is sufficiently small compared to the radius of the spreading population.

### 2.3 Data

In this study we use synthetic data generated by the RWMs described in Section 2.2 to produce count data with properties analogous to those of relevant experiments. The work is presented as two different cases: (i) Case 1 uses data generated from a non-interacting RWM described in Section 2.2.1; and, (ii) Case 2 uses data generated from an interacting RWM described in Section 2.2.2. For Case 2, we explore results based on data collected at the end of two identically prepared simulations: Case 2a uses early-time data after 100 time steps and Case 2b uses late-time count data after 600 time steps. As we demonstrate, this distinction has an important impact on parameter identifiability as we will detail in Section 3.2. In all cases our data are stored in a vector denoted *y*° noting that in the applied statistics literature it is customary not to use bold face notation for vectors.

In reality, generating count data from high-density experiments is very time-consuming, and this can be addressed by subsampling the data. We take the same approach by working with subsampled count data from our RWMs to ensure that our methods do not rely on working with impractical amounts of cell counts. To this end, we collect *N* data points for each subpopulation from the full count data. Consequently, *y*° is a vector of length 2*N* for Case 1, whereas *y*° is a vector of length *N* for Case 2.

In Case 1, we simulate the non-interacting RWM on a 200 *×* 20 lattice with (*P, ρ, λ*_1_, *λ*_2_)^*⊤*^ = (1, 0.5, 0.02, 0.03)^*⊤*^. Initially, red and green agents occupy sites within columns 81 *≤ i ≤* 120, with relatively low initial densities of 0.5 and 0.2, respectively, as in Figure 2(a). Full count data at *t* = 0 for the red and green subpopulations are shown in Figure 2(c)–(d). Data are then collected after *k* = 100 time steps, producing the snapshots in Figure 2(b), where we can see that the total population has spread, drifted in the positive *x*-direction, and the total number of agents grows over the simulation. Full count data for both the red and green subpopulations after *k* = 100 time steps are presented in Figure 2(e)–(f), respectively. We subsample *N* = 12 equally-spaced data points from the full count data in Figure 2(e)–(f) for both subpopulations, given in Figure 2(g)–(h). We denote the sampled data as 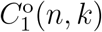 and 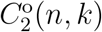, where *n* = 1, 2, 3, …, *N* and *k* = *K* = 100. The relationship between the data index *n* and the lattice column index *i* is *i* = (*n* − 1)18.

**Figure 2.**
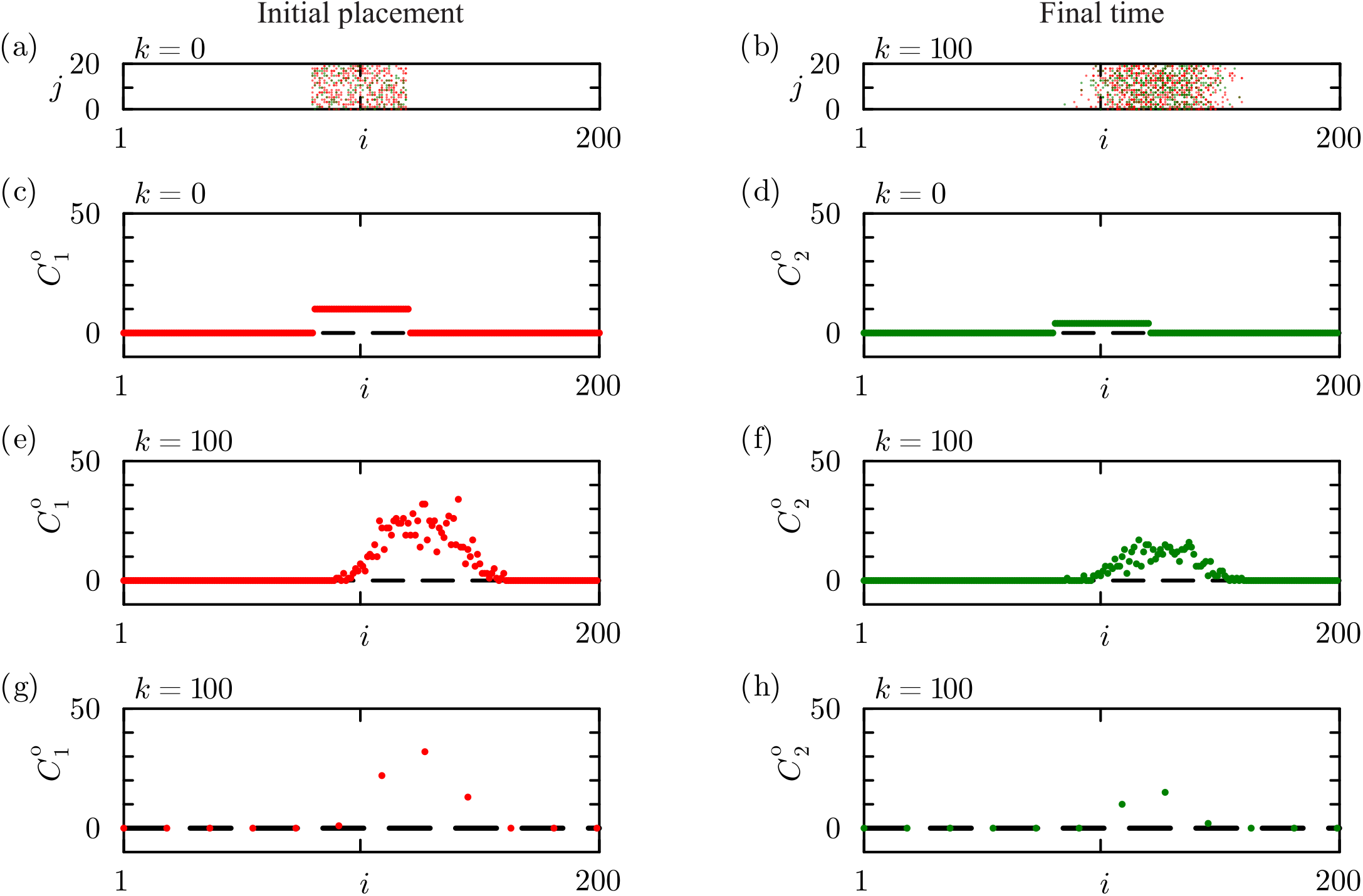
Snapshots of agent distributions and count data from the non-interacting RWM with (*P, ρ, λ*_1_, *λ*_2_)^*⊤*^ = (1, 0.5, 0.02, 0.03)^*⊤*^. (a)–(b) Snapshots of agent distribution. (c)–(f) Full count data. (g)–(h) Subsampled count data obtained from data in (e)–(f), respectively. Horizontal dashed lines in (e)–(h) show the 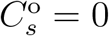 lower bound for the count data. Results correspond to *I* = 200 and *J* = 20.

In Case 2, we simulate the interacting RWM on a 200 *×* 200 lattice with (*P, λ, c*_0_)^*⊤*^ = (1, 0.01, 0.8)^*⊤*^. Agents are initially placed within a circular region of radius *r*_0_ = 20Δ. Within this region, each site is occupied by a single agent with probability 0.8 to reflect the relatively high density design of the barrier assay. A snapshot of the initial condition is shown in Figure 3(a). The early-time snapshot after *k* = 100 time steps, and a late-time snapshot after *k* = 600 time steps are shown in Figure 3(b)–(c), respectively. A green rectangular transect is superimposed on each snapshot, with detailed views of the corresponding transects shown in Figure 3(d)–(f), respectively. The corresponding full count data from the rectangular transect after *k* = 0, 100 and 600 time steps are shown in Figure 3(g)–(i), respectively. We subsample *N* = 20 equally-spaced data points from the full count data after *k* = 0, 100 and 600 time steps. The subsampled data is presented in Figure 3(j)–(l), respectively. In Case 2, we denote the sampled data as *C*°(*n, k*), with *k* = 100 for Case 2a and *k* = 600 for Case 2b. The relationship between the data index *n* and the column index *i* in Case 2 is given by *i* = (*n* − 1)10.

**Figure 3.**
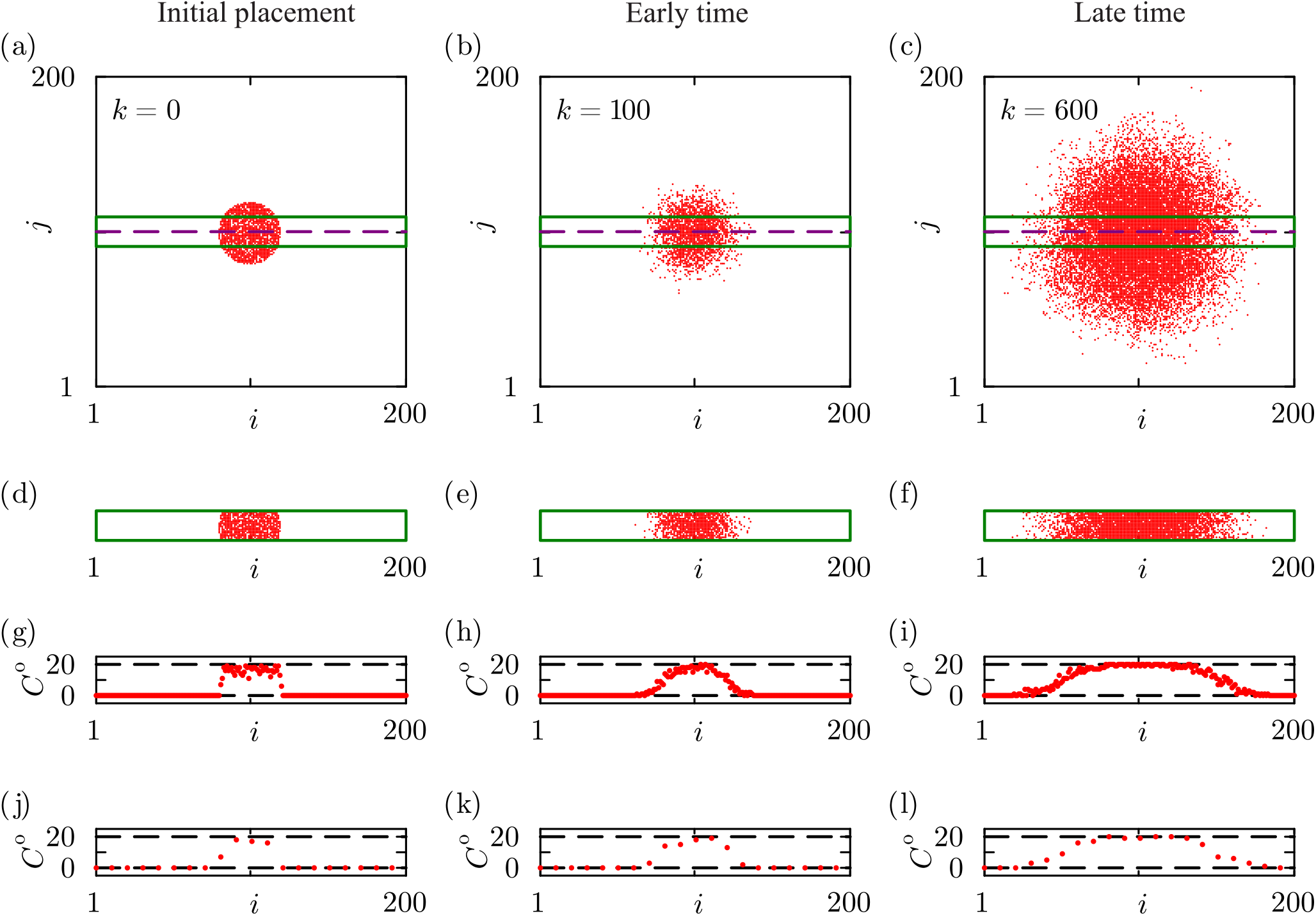
Snapshots of agent distributions and count data from the interacting RWM with (*P, λ, c*_0_)^*⊤*^ = (1, 0.01, 0.8)^*⊤*^. (a)–(c) Snapshots of agent distribution superimposed with a green rectangular transect of height *H* = 20 and purple line showing the centre of the spreading population where *j* = 100. (d)–(f) Agent distributions within the transects in (a)–(c), respectively. (g)–(i) Count data from the snapshots in (d)–(f), respectively. (j)–(l) Subsampled count data obtained from data in (g)–(i), respectively. Horizontal dashed lines in (g)–(l) show the *C*° = 0 lower bound and *C*° = 20 upper bound for the count data. Results correspond to *I* = *J* = 200 and *H* = 20.

In this work Cases 2a and 2b illustrate that estimation, identifiability, and sensitivity analysis can be performed across varying experimental timescales. As we will demonstrate, a key feature of Case 2 is that the identifiability of various simulation parameters depends upon the simulation timescale. For completeness, we also repeat a similar analysis using late-time data for Case 1 in Section 4 of the Supplementary Material. Further exploration involving different choices of parameter values and simulation durations can be explored by adapting open-access software on GitHub.

### 2.4 Likelihood function

In this section, we describe how two physically-motivated measurement error models that probabilistically relate the count data summarised in Figure 2–3 to appropriate solutions of the relevant surrogate PDE models. For Case 1, we use a Poisson-based measurement error model, whereas for Case 2 we use a binomial measurement error model. These different choices of measurement error models accommodate differences in the properties of the count data, which we now describe.

For the red subpopulation in Case 1, we interpret the integer count data 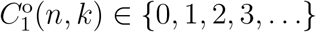 at a given spatial location after *k* time steps as representing a finite number of independent samples from an underlying stochastic process (i.e. the RWM or a real biological experiment). These data are considered to be samples from a distribution in which the expected total number of agents in the column corresponding to the *n*th data point is related to the solution of the continuum-limit model, given by *J × c*_1_(*x*_*n*_, *t*_*k*_). Here, *x*_*n*_ denotes the horizontal Cartesian coordinate associated with the *n*th data point and *t*_*k*_ is time after *k* time steps. This relationship, together with the non-negative, unbounded integer count data motivates the use of a Poisson measurement error model to link the observed count data with the continuum-limit solution. While other options are possible, we use a Poisson-based measurement error model for simplicity as it does not introduce any additional parameters to be estimated [13]. Following a similar argument we can link count data from the green subpopulation to the solution of Equation (2)–(3) using a Poisson measurement error model. Invoking a standard independence assumption, we can evaluate the probability density for each subpopulation and take the product of these two probability densities, which yields a Poisson-based likelihood for the non-interacting RWM in Case 1 at a single spatial location *x*_*n*_:

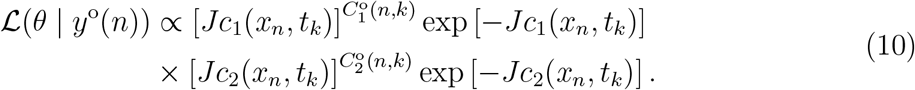

Invoking a standard independence assumption we can consider count data at all spatial locations to give a log-likelihood function as follows,

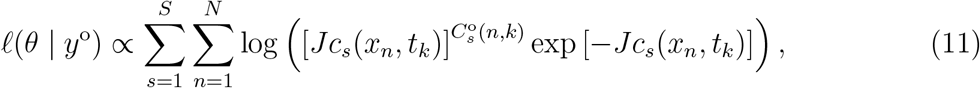

where *S* = 2 in for our RWM.

Count data for Case 2 consists of agent counts within each column of the transect, which implicitly determine the number of vacant lattice sites in each column. Thus, the data take the form *C*°(*n, k*) and *E*°(*n, k*) = *H* − *C*°(*n, k*) for *n* = 1, 2, 3, …, *N*. Interpreting *C*°(*n, k*) at a given spatial location as a finite number of independent samples from an underlying stochastic process (i.e. the RWM or a real biological experiment), these samples approximate a continuous measurement of agent occupancy within that column. These data are effectively drawn from a distribution in which the expected occupancy fraction is given by the solution of the continuum-limit model, *H × c*(*r, t*). However, while *C*°(*n, k*) represents count data along the centreline of the spreading population, *H × c*(*r, t*) denotes the counts along the radius. By symmetry, we have *r* = |*x* − *x*_0_|, where *x*_0_ denotes the *x*-coordinate of the centre of the spreading population. These observations, together with the fact that the count data are bounded non-negative integers, allow us to make use of a binomial measurement error model to link the observed count data with the solution of the PDE model, as the binomial distribution is defined over non-negative integers with a finite upper bound [13]. Here, we adopt a binomial measurement error model for Case 2 at a single spatial location *x*_*n*_ leading to a likelihood function of the form

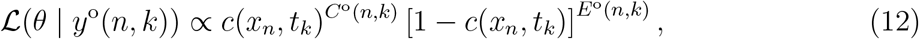

giving rise to the log-likelihood function for data across multiple spatial locations as

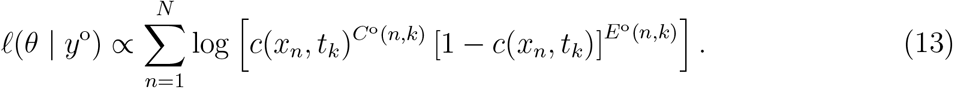

In this work, we set the proportionality constants in Equation (11) and Equation (13) to unity [41].

### 2.5 Likelihood based estimation and identifiability

In all cases we aim to estimate parameters in terms of rates within the stochastic model. For example, we will estimate 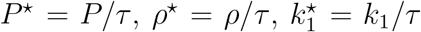 and 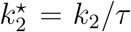 for Case 1. For a set of count data *y*° together with a relevant process and measurement error model, we have access to a log-likelihood function, *ℓ*(*θ* | *y*°). The parameters that best match the data, 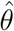, correspond to the choice of *θ* that maximises *ℓ*(*θ* | *y*°), giving rise to the maximum likelihood estimate (MLE):

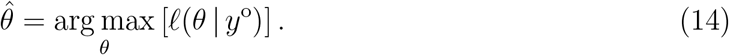

Throughout this work we use the Nelder–Mead algorithm with simple bound constraints [67] to compute 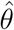 [68]. In general, our numerical optimisation calculations for identifiable parameters are robust to the choice of initial estimates for the iterative solver, and the algorithm converges using the default stopping criteria.

To compare results against asymptotic thresholds, we work with a normalised log-likelihood function

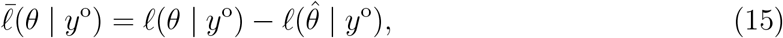

so that 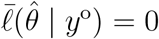. Given the normalised log-likelihood function, we use the profile likelihood approach to quantify the precision of parameter estimates by examining the curvature of the log-likelihood function [35, 37–40]. We partition the full parameter set *θ* into interest parameters, *ψ*, and nuisance parameters, *ω*, such that *θ* = (*ψ, ω*)^*⊤*^. In this work, we focus exclusively on univariate profile log-likelihood functions, where the interest parameter is always a single parameter. For a given dataset *y*°, the profile likelihood for the interest parameter *ψ* under the partition *θ* = (*ψ, ω*)^*⊤*^ is defined as:

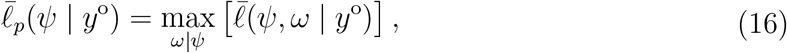

which implicitly defines a function *ω*^*†*^(*ψ*), representing the optimal values of *ω* for each value of *ψ*. Similar to the MLE, we calculate all profile likelihoods using the Nelder-–Meadalgorithm using the same bound constraints and default stopping criteria. As a specific example, consider the non-interacting RWM in Case 1 with 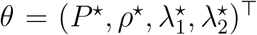 where we can compute four univariate profile likelihood functions by choosing the interest and nuisance parameters as follows: (i) *ψ* = *P* ^*⋆*^ with 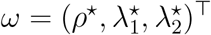; (ii) *ψ* = *ρ*^*⋆*^ with 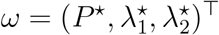; (iii) 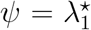 with 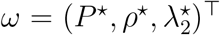; and, (iv) 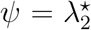 with 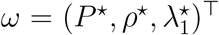. For all profile likelihood calculations, we evaluate 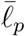 over a uniformly-discretised interval containing the MLE. For example, for *ψ* = *P* ^*⋆*^ we consider the interval 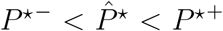, and compute 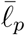 across a uniform discretisation of that interval. The resulting univariate profile likelihood function can be visualised by simply plotting 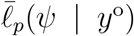. The curvature of 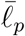 provides information about the practical identifiability of the interest parameter [37]. A flat profile likelihood indicates that the parameter is not identifiable [34], whereas the degree of curvature of 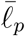 indicates inferential precision [35, 37–40]. We determine likelihood-based confidence intervals by identifying intervals where 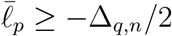, where Δ_*q,n*_ denotes the *q*th quantile of the *χ*^2^ distribution with *n* degrees of freedom, which we take to be the relevant number of free parameters [69, 70]. For example, identifying the interval where 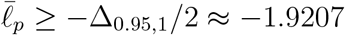 allows us to compute the asymptotic 95% confidence interval [4, 37] for a univariate profile likelihood [41].

### 2.6 Parameter-wise prediction

To propagate uncertainty in a single interest parameter *ψ* into the uncertainty in the model solution, we define an interval for *ψ* such that 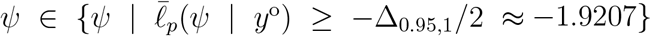 [57]. We take a uniform discretisation of that interval, with *M* mesh points, and solve the model at each mesh point with 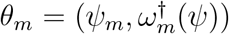 for *m* = 1, 2, 3, …, *M*. For each of these *M* model solutions, we construct a prediction interval for data realisations [4] as follows. Each solution of the surrogate model can be interpreted as the mean of the relevant noise model, and we account for the width of the noise model distribution by computing the 5% and 95% quantiles of the measurement error model at each spatial location. The union of these *M* intervals at each spatial location yields a parameter-wise prediction interval for data realisations. In this work, we use *M* = 100 and the procedure can be repeated for all parameters in *θ*.

As a concrete example, consider the non-interacting RWM with 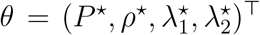. To construct parameter-wise prediction interval for *ψ* = *P* with 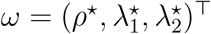, we first define the interval 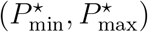. As we will illustrate, for identifiable problems the bounds of this interval are values of *P* ^*⋆*^ for which 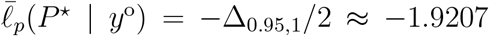. These bounds can be determined using a root finding method (e.g. we use the bisection algorithm). For non-identifiable parameters, a sufficiently flat univariate profile likelihood function can fail to intersect one or both asymptotic thresholds. In such cases, user-defined bounds can be implemented, as we will illustrate in Section 3.2. Once the interval 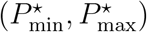 is identified, it is uniformly-discretised into *M* mesh points, giving 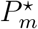 for *m* = 1, 2, 3, …, *M*. For each 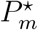, we compute the corresponding nuisance parameters 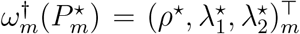 and solve the surrogate PDE for the *M* parameter combinations to give 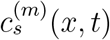 for *m* = 1, 2, 3, …, *M* and *s* = 1, 2, …, *S*. Each PDE solution is uniformly-discretised over space, giving 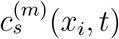, for *i* = 1, 2, 3, …, 200. For each of these discretised solutions, we define a prediction interval for the data realisation as 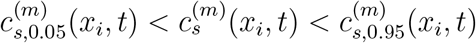, where the lower and upper bounds correspond to the 5% and 95% quantiles of the Poisson-based measurement error model, respectively [41]. After calculating these intervals at each spatial location for the *M* solutions, we take the union of the *M* solutions at each spatial location to give the parameter-wise prediction interval at the *i*th spatial location

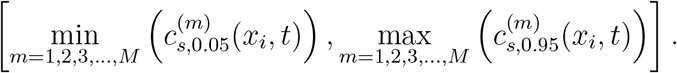

A similar procedure is applied to the interacting RWM in Case 2 to construct various parameter-wise prediction intervals. This computational approach provides a complete representation of the parameter-wise prediction intervals, which can be interpreted as a form of tolerance interval [71]. These intervals account for both individual parameter uncertainty and data realisation uncertainty in an efficient, objective manner that avoids random sampling such as in a more standard LHS-PRCC approach [41].

## 3 Results and Discussion

In this section, we present the results for the estimation, identifiability analysis, and sensitivity analysis using the data described in Section 2.3 for both the non-interacting and interacting RWMs. For the non-interacting RWM in Case 1, we present one set of results corresponding to a simulation at early time, the result corresponding to a late-time simulation is presented in Section 4 of the Supplementary Material document. For the interacting RWM in Case 2, we present two results: Case 2a corresponds to an early-time simulation, and Case 2b to a late-time simulation. The results are presented in Figure 4–6, where each row corresponds to a different parameter, and each column (from left to right) shows the univariate profile likelihood, the parameter-wise prediction interval, and the difference between the confidence set and the solution of the continuum-limit evaluated at the MLE, respectively.

**Figure 4.**
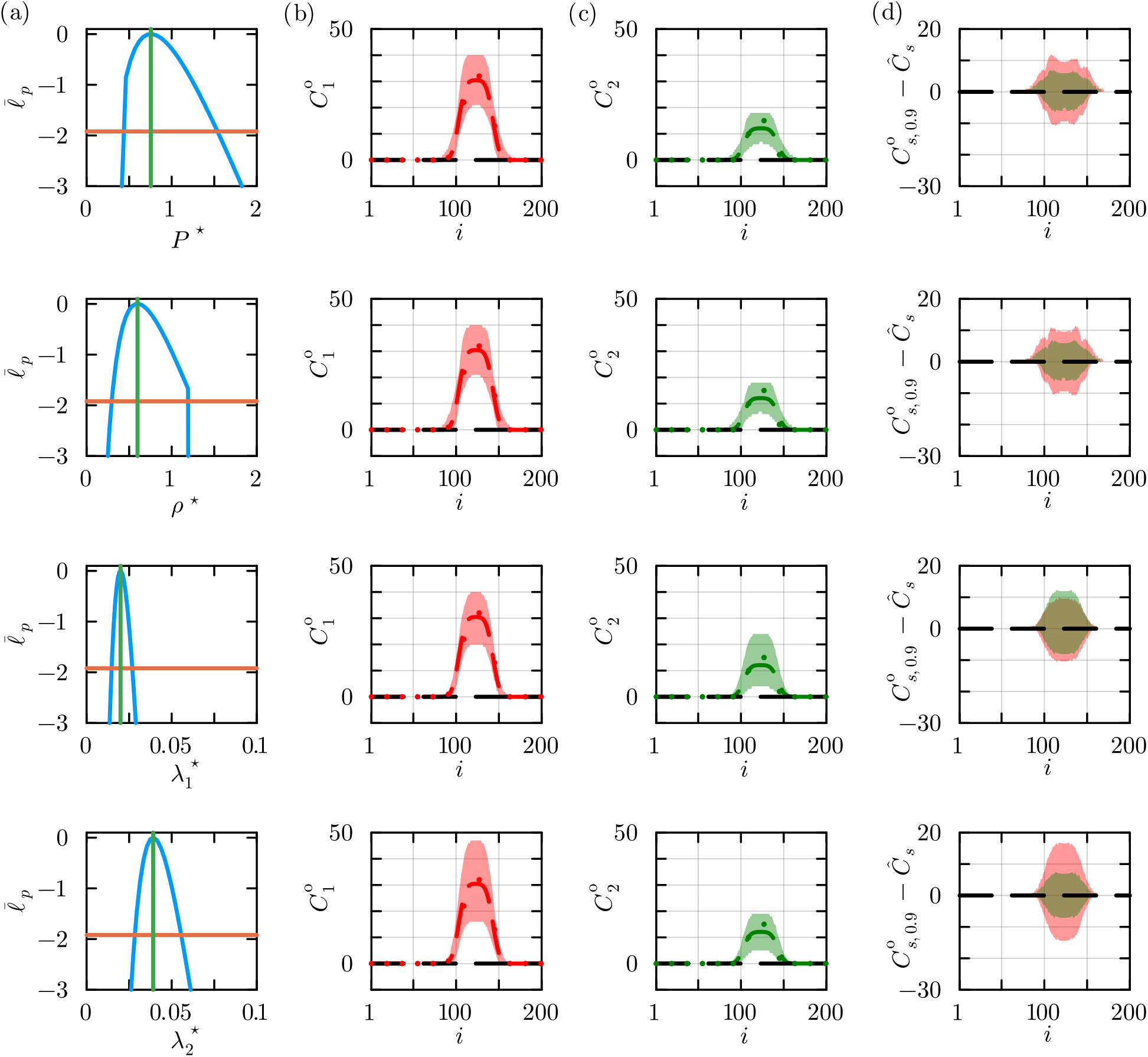
Estimation, identifiability and sensitivity results for the non-interacting RWM. (a) Univariate profiles for 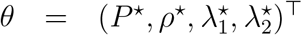 indicate the MLE (vertical green line) and the 95% asymptotic threshold (horizontal orange line). The MLE is 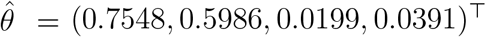. (b)–(c) Profile-wise prediction intervals for *C*_1_ (shaded red) and *C*_2_ (shaded green) for each parameter superimposed with count data (green dots), and the solution of the PDE evaluated at the MLE for *C*_1_ (dashed red) and *C*_2_ (dashed green). (d) Difference between confidence set and the solution of the mathematical model evaluated at the MLE for *C*_1_ (shaded red) and *C*_2_ (shaded green).

### 3.1 Case 1

Using the noisy count data collected from the non-interacting RWM at early-time, presented in Figure 2(g)–(h), we use the Poisson-based measurement error model and associated log-likelihood function to estimate the MLE, 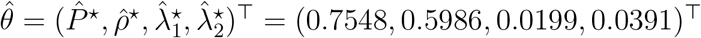. Compared with the true values, 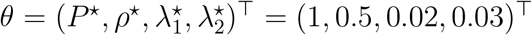, these point estimates appear quite reasonable but they do not provide any indication of their identifiability. We explore the practical identifiability of each individual parameter by constructing various univariate profile likelihoods shown in column (a) of Figure 4. Each profile likelihood is relatively narrow with a single peak around the MLE which indicates that all parameters are practically identifiable. For each parameter, we define a 95% asymptotic confidence interval as the region where 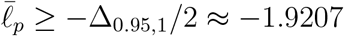 [69]. For example, to construct the univariate profile likelihood for the interest parameter *P* ^*⋆*^, as shown in the first row of column (a) in Figure 4, we evaluate 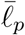 over a specified interval 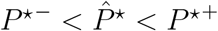, where we choose *P* ^*⋆*−^ = 0 and *P* ^*⋆*+^ = 2. The 95% confidence interval for *P* ^*⋆*^ is computed using the bisection algorithm, giving 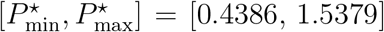. The 95% confidence intervals for the remaining parameters in Figure 4 are: (i) *ρ*^*⋆*^ = 0.5986 [0.2961, 1.1925]; (ii) 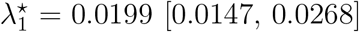, and (iii) 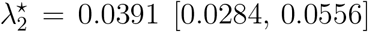. The true value for each parameter lies within the corresponding 95% confidence interval, indicating that the estimates are reasonably precise. The relatively narrow shape of each univariate profile provides further evidence that all parameters are well identified by the count data we have considered.

Parameter-wise sensitivity analysis is performed by constructing parameter-wise prediction intervals, as shown in columns (b)–(c) of Figure 4. Each sub-figure in these columns illustrates how the results at early-time simulation depend on each of the parameters, with results in column (b) detailing the impact of a single parameter on 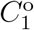 and results in column (c) showing the impact on 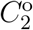. In column (d) of Figure 4 we superimpose plots of the difference between the respective confidence sets for model realisations and the surrogate PDE solution evaluated at the MLE using the same visualisation approach as in [4]. Here 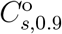 is the confidence set bounded by 5% and 95% quantiles of the relevant distribution associated with the measurement error model. These plots effectively show the width of the parameter-wise prediction interval for each parameter as a function of position. This analysis provides detailed insight into the width of the parameter-wise prediction intervals across spatial locations and enables a comparison between the parameter sensitivity for 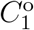 and 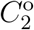.

All parameter-wise prediction intervals in Figure 4 are wider in the middle regions of the population (i.e. 100 ⪅ *i* ⪅ 150), indicating larger sensitivity in the predicted count data in the bulk of the population, and a smaller sensitivity in the tail regions of the population (i.e. *i* ⪅ 100 and *i* ⪆ 150). This pattern is consistent with our intuitive expectations, as the higher density middle regions can involve larger stochastic variations compared to the low or zero agent density in the tail region characterized by minimal variations. The parameter-wise prediction interval for 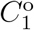 associated with 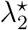, shown in the last row of column (b) in Figure 4 is noticeably wider than the prediction intervals for the other parameters, thus suggesting that 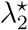 has a stronger influence on 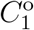 than the other parameters. Additionally, 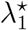 appears to affect both 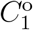 and 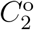 similarly, while 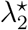 has a larger impact on 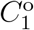 and a reduced impact on 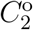.

All results in Figure 4 are based on early-time (*k* = 100) count data, and the associated profile likelihoods indicate that all four parameters are identifiable. To assess whether these main findings are sensitive to the timing of data collection, we conduct a similar analysis using count data collected at a later time point, as reported in Section 4 of the Supplementary

Material. In brief, we find that the conclusions remain consistent regardless of whether we consider early or later-time data. Interestingly, this is different from the case for the interacting RWM in Case 2 as we will now explore and discuss.

### 3.2 Case 2

Using noisy count data from the interacting RWM at early-time, presented in Figure 3(k) for Case 2a, we aim to estimate: (i) the motility rate *P* ^*⋆*^; (ii) the proliferation rate *λ*^*⋆*^; and (iii) the initial density *c*_0_. In Case 2, we include the initial condition *c*_0_ as an unknown parameter since the initial number of cells in a barrier assay is not always known precisely [23]. Using the log-likelihood function based on the binomial measurement error model, the MLE is 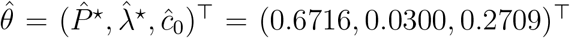. These estimates differ substantially from the true values, *θ* = (*P* ^*⋆*^, *λ*^*⋆*^, *c*_0_)^*⊤*^ = (1, 0.01, 0.8)^*⊤*^. The large discrepancies arising from the use of noisy data suggest a potential indication of challenges with parameter identifiability that we will now explore.

We explore the practical identifiability of each parameter for Case 2a by constructing three univariate profile likelihoods shown in Figure 5, column (a). The profiles likelihood for both *P* ^*⋆*^ and *c*_0_ exhibit a single peak around the MLE, but both profiles are very wide indicating that these parameters are not well-identified by this data, with 95% confidence intervals of *P* ^*⋆*^ = 0.6716 [0.3718, 1.3822] and *c*_0_ = 0.2709 [0.0263, 0.8820], respectively. The profile likelihood for *λ*^*⋆*^ indicates that *λ*^*⋆*^ is one-sided identifiable, with the 95% asymptotic confidence interval bounded below at *λ*^*⋆*^ = 0.0097. We construct the profile likelihood for *λ*^*⋆*^ using a conservative upper bound of 0.04. Values of *λ*^*⋆*^ *>* 0.04 are not considered, as simulations with *λ*^*⋆*^ = 0.04 show a clear mismatch with the data, indicating that larger values are not biologically relevant. The poorly identified initial density *c*_0_ suggests that the data collected after *k* = 100 time steps are insufficient to accurately identify *c*_0_. Since the data are collected relatively early in the simulation (i.e. the number of time steps is small compared to the proliferation time scale 1*/λ*), the simulation does not involve significant agent proliferation, which explains why *λ*^*⋆*^ is poorly identified. These results highlight that parameter non-identifiability can arise even in the relatively simple setting of Case 2a, in contrast to Case 1, where all parameters are identifiable.

**Figure 5.**
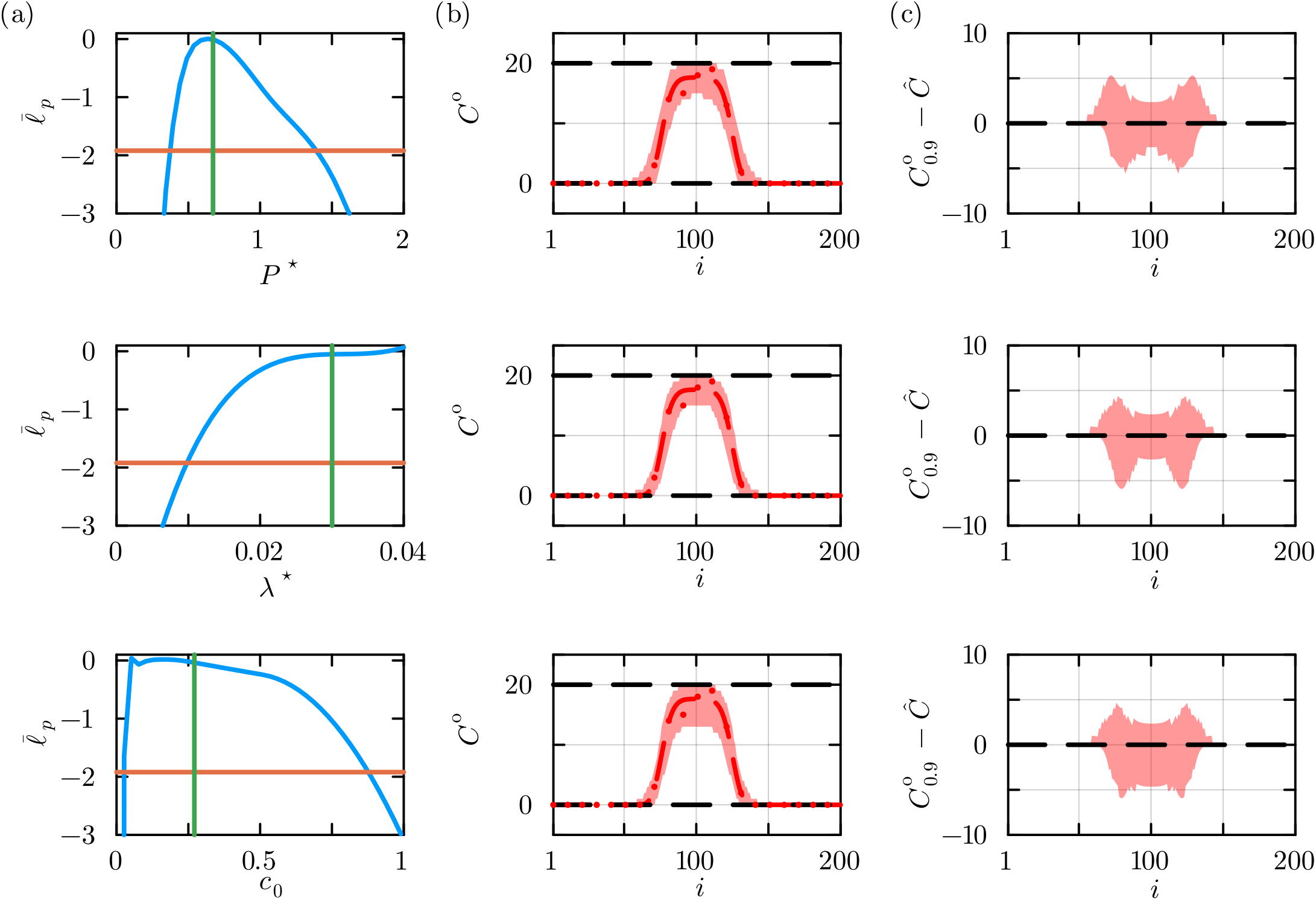
Estimation, identifiability and sensitivity results for the non-interacting RWM using early-time data. (a) Univariate profiles for *θ* = (*P* ^*⋆*^, *λ*^*⋆*^, *c*_0_)^*⊤*^ indicate the MLE (vertical green line) and the 95% asymptotic threshold (orange horizontal line). The MLE is 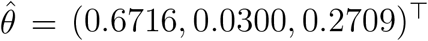. (b) Profile-wise prediction intervals for *C* (shaded red) for each parameter superimposed with count data (red dots) and the solution of the PDE evaluated at the MLE. (c) Difference between confidence set and the solution of the mathematical model evaluated at the MLE.

Sensitivity analysis for Case 2a is performed by constructing parameter-wise prediction intervals shown in Figure 5, column (b). We use *λ*^*⋆*^ = 0.04 as the upper bound when constructing the parameter-wise prediction intervals for the non-identifiable parameter *λ*^*⋆*^, as 0.04 is the maximum plausible *λ*^*⋆*^ value considered in the model. Each sub-figure in Figure 5, column (b), illustrates how the early-time simulation results depend on each parameter: *P* ^*⋆*^, *λ*^*⋆*^, or *c*_0_, respectively. We find no obvious differences in the parameter-wise prediction intervals for these three parameters, indicating that all three parameters have a similar impact on the model outcome. The differences between the respective confidence sets and the PDE solution evaluated at the MLE for each parameter in column (b) of Figure 5 show that the prediction intervals in Case 2a are relatively narrow in the middle region of the spreading population (i.e. 80 ⪅ *i* ⪅ 120) and wider in the transition regions (i.e. 70 ⪅ *i* ⪅ 80 and 120 ⪅ *i* ⪅ 130). This pattern is in contrast to Case 1, where the middle region of the population exhibited larger variability. The differences are because of the crowding mechanism of RWM in Case 2, which ensures count data are bounded above by *H* = 20. In the high-density middle region, fluctuations are suppressed as the agent count approaches *H*, thereby reducing variability. In contrast, the non-interacting RWM in Case 1 has no such upper bound, allowing increasing fluctuations in high-density regions. The relatively wider prediction intervals in the transition regions of Case 2a are explained by the fact that fluctuations in count data in these regions do not reach the upper or lower bounds, leading to greater variability compared to the middle region. Both observations are also supported by the data shown in Figure 3(e) and (h).

In Case 2b, we perform a similar analysis using data from late-time simulation (*k* = 600), as presented in Figure 3(l). Using the same binomial measurement error model, the MLE for Case 2b is given by 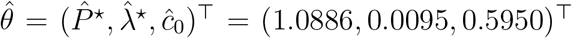. The estimates of *P* ^*⋆*^ and *λ*^*⋆*^ are close to the expected true values (*P* ^*⋆*^ = 1, *λ*^*⋆*^ = 0.01), while the estimate for *c*_0_ differs more substantially from the expected true value (*c*_0_ = 0.8). The univariate profile likelihoods presented in the first and second rows of Figure 6, column (a), show that *P* ^*⋆*^ and *λ*^*⋆*^ are well identified by this data, with relatively narrow profiles and 95% confidence intervals of *P* ^*⋆*^ = 1.0886 [0.8531, 1.4201] and *λ*^*⋆*^ = 0.0095 [0.0077, 0.0119]. The flat profile likelihood for *c*_0_ indicates that this parameter is not well-identified by this data. This finding differs from Case 2a, where *λ*^*⋆*^ is non-identifiable and *c*_0_ is identifiable. This difference in identifiability reflects the timing of data collection. In Case 2b, the data are collected at a late-time simulation (*k* = 600), by which time the system has evolved for a sufficient duration of time that the influence of the initial condition has diminished. Interestingly, the extended simulation time allows for agent proliferation to have a more pronounced impact which means that *λ*^*⋆*^ is well-identified by this late-time data.

**Figure 6.**
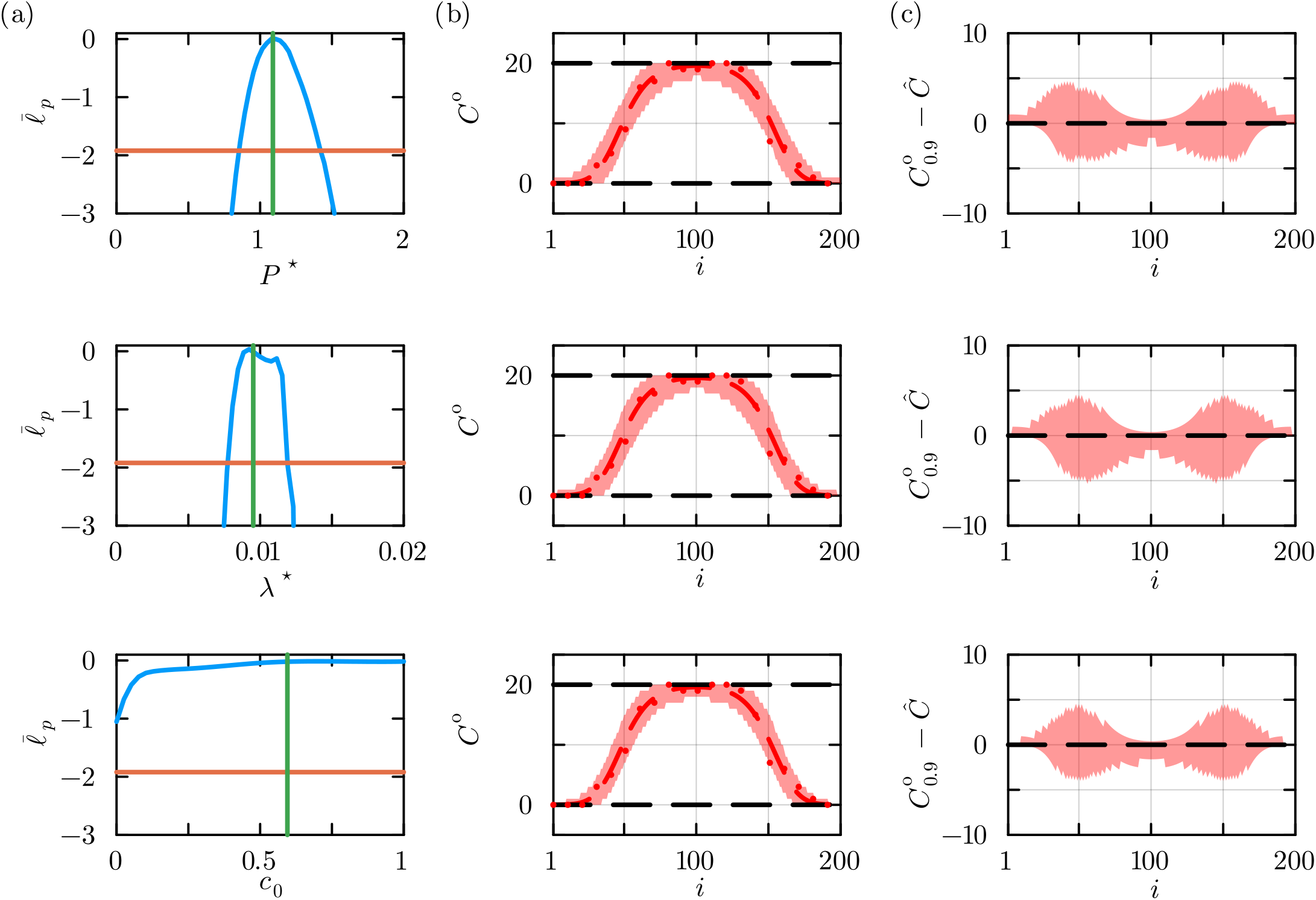
Estimation, identifiability and sensitivity results for the non-interacting RWM using late-time data. (a) Univariate profiles for *θ* = (*P* ^*⋆*^, *λ*^*⋆*^, *c*_0_)^*⊤*^ indicate the MLE (vertical green line) and the 95% asymptotic threshold (orange horizontal line). The MLE is 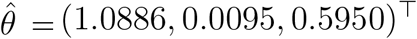. (b) Profile-wise prediction intervals for *C* (shaded red) for each parameter superimposed with count data (red dots) and the solution of the PDE evaluated at the MLE. (c) Difference between confidence set and the solution of the mathematical model evaluated at the MLE.

Sensitivity analysis for Case 2b involves constructing various parameter-wise prediction intervals shown in Figure 6, column (b). For the non-identifiable parameter *c*_0_, the prediction interval is constructed over the range of all physically–relevant values, *c*_0_ ∈ [0, 1]. As with Case 2a, we see no obvious differences in the parameter-wise prediction intervals for these three parameters. Plotting the differences between the boundary of the confidence sets and the continuum model solution evaluated at the MLE, given in Figure 6 column (c), we see that the width of the prediction interval in the middle region of the population is even narrower than in Case 2a. This occurs because the data in Case 2b are collected at a later simulation time, during which the agent density in the middle region continues to increase towards the carrying capacity density, as shown in Figure 3(f) and (i). At these higher densities, the fluctuations are further suppressed, leading to reduced variability. In fact, agent counts in the middle region of the population reach the upper bound *H* = 20, which means that there are no fluctuations above this value owing to crowding effects in the RWM. This explains why the width of the prediction interval above the MLE becomes zero in the centre of the middle region of the spreading population.

## 4 Conclusion and Future Work

In this work we present a novel and efficient approach for sensitivity analysis that can be applied to computationally expensive RWMs. A key element of our approach is to link sparse noisy count data to the solution of an efficient surrogate PDE model using various forms of physically-motivated measurement error models. These tools place our analysis within a likelihood-based framework that can be used for parameter estimation, parameter identifiability analysis and sensitivity analysis via the targeting property of the profile likelihood. Count data from a classical non-interacting RWM are described using a Poisson-based measurement error model, while count data from an interacting exclusion process-based RWM is captured using a binomial measurement error model. Within this general framework we illustrate how to obtain point estimates using maximum likelihood estimation, we analyse the identifiability of those estimates using profile likelihood, and we use parameter-wise predictions as a form of sensitivity analysis. Across our two case studies, we illustrate how the approach applies to both identifiable and poorly-identified problems, and we illustrate how the approach can be implemented using exact or numerical solutions of the relevant surrogate PDE.

Our approach makes use of the targeting property of the profile likelihood to focus on parameter estimates that are targeted towards those regions in the parameter space where the solution of the surrogate model is close to the data. This is a significant departure from random sampling-based approaches, such as the LHS-PRCC approach, that relies on random parameter samples to explore parameter sensitivity. A key limitation of working with random parameter samples is that those samples are not necessarily guided by the aim of keeping the model solution close to the target data. While all parameter-wise prediction analysis implemented in this work focuses on univariate parameter-wise prediction intervals, it is possible to use the same approach to generate multivariate prediction intervals (e.g. bivariate profile-wise predictions [72]) where the interest parameters are taken to be a small group of model parameters instead of a single parameter.

Our methodology for sensitivity analysis is demonstrated via two biologically-motivated case studies. Each case study involves a particular choice of process model in the form of a computationally expensive lattice-based RWM and a physically-motivated measurement error model that is guided by the properties of the data generated by the RWM. While we focus on making particular choices of biologically-motivated process and measurement error models, it is important to point out that this framework can be applied quite generally by varying either the process model, the measurement error model or both the process and measurement error models. This means that the same framework can be applied to many extensions of the current case studies. For example, in Case 1 we assume that both subpopulations of agents share the same migration parameters *P* and *ρ*. This assumption can be easily relaxed to allow each subpopulation to have unique parameters so that *θ* = (*P*_1_, *P*_2_, *ρ*_1_, *ρ*_2_, *λ*_1_, *λ*_2_)^*⊤*^, where *P*_*i*_ and *ρ*_*i*_ represent the motility probabilities and the bias parameter for the *i*th subpopulation, respectively. Under this extension the relevant surrogate PDE model is given by

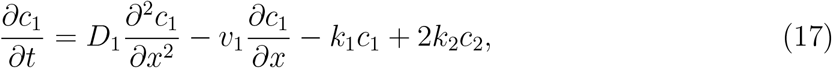

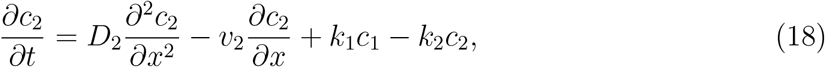

With

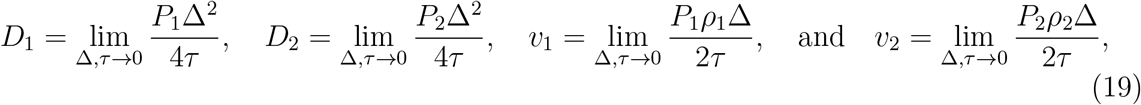

where *k*_1_, and *k*_2_ are defined previously in Equation (4). For this extension the surrogate model is best solved numerically rather than generalising the transform approach outlined in the Supplementary Material document. The rest of the workflow for estimation, identifiability and parameter-wise sensitivity analysis remains unchanged.

In this work we use a Poisson-based measurement error model for data generated by the non-interacting RWM because the Poisson distribution is defined over unbounded, non-negative integers. We note that other distributions, such as the negative binomial distribution, also share these properties and could serve as alternative measurement error models [73]. Similarly, we use a binomial measurement error model for data generated by the interacting RWM since the binomial distribution is defined over non-negative integers with a finite upper bound. The truncated Poisson distribution has similar features and so could be used as an alternative measurement error model [74].

All results in our case studies are presented for data generated at a single time point only. While many experiments focus only on data at a single time point, it is conceptually and computationally straightforward to generalise our approach to apply to problems where data is collected at *K* time points for *S* subpopulations [41]. For example, a Poisson-based log-likelihood function with count data collected at *K* time points across *S* subpopulations can be written as

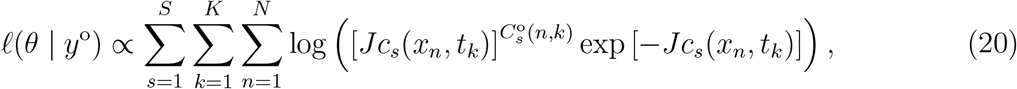

where the data vector *y*° has length *S × K × N*. With this modest generalisation of the log-likelihood function, all other steps for estimation, identifiability analysis, and parameter-wise sensitivity analysis remain unchanged. A similar generalisation can be made for the log-likelihood function based on the binomial measurement error model with data collected at *K* time points across *S* subpopulations, which can be expressed as

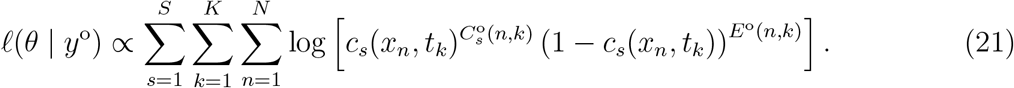

Both Equation (20) and Equation (21) can be further explored in the context of more complex models.

A key feature of our work is to use approximate continuum-limit PDE models as computationally-efficient surrogates. While in many cases the continuum-limit models are well–established in the literature, and known to lead to accurate approximations [15, 19], it is not strictly necessary to use this approach. Other methods for deriving approximate surrogate models from data, such as working with Gaussian processes [75, 76] or equation learning methods can also be used [77, 78], and we leave these approaches for future exploration.

## SUPPLEMENTARY MATERIAL

## S1 Exact solution of surrogate PDE for Case 1

Here, we implement a transformation method to obtain analytical solutions for systems of linear partial differential equations (PDEs) with coupled source terms relevant to Case 1. This method, based on diagonalisation, can be applied to systems with *n* subpopulations [1] such as

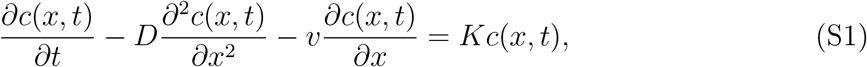

where *c*(*x, t*) is a vector of length *n, K* is an *n × n* constant matrix, and *D* and *v* are scalar constants. As we will illustrate, a key assumption in this approach is that the matrix *K* is diagonalisable.

The first step in the solution procedure is to determine an *n × n* constant matrix *S*, where each column of *S* corresponds to an eigenvector of *K* [1]. We then define a second vector of length *n*, given by *b*(*x, t*) = *S*^−1^*c*(*x, t*). Consequently, we can write *c*(*x, t*) = *Sb*(*x, t*), and Equation (S1) can be expressed as:

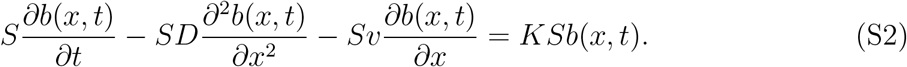

Left-multiplying Equation (S2) on both sides by *S*^−1^ yields:

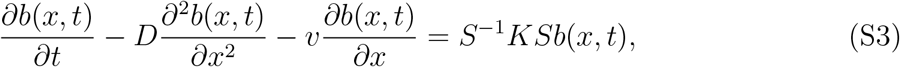

where *S*^−1^*KS* is a constant diagonal matrix [1]. Equation (S3) thus decouples into *n* independent PDEs, each of which can be solved analytically for certain choices of boundary conditions and initial conditions. Once *b*(*x, t*) is determined, the solution is given by *c*(*x, t*) = *Sb*(*x, t*).

Now consider the system of PDEs, Equations(2)–(3) in the main document, with *c*(*x, t*) = (*c*_1_(*x, t*), *c*_2_(*x, t*))^*⊤*^,

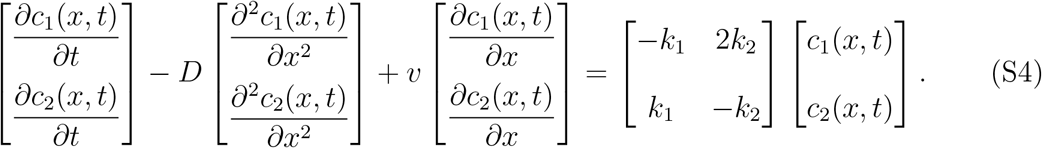

We consider this PDE model on an infinite domain −*∞ < x < ∞*, with the initial condition for *c*_1_(*x, t*) given by *c*_1_(*x*, 0) = *α >* 0 for |*x* − *x*_0_| *≤ h*, and *c*_1_(*x*, 0) = 0 for |*x* − *x*_0_| *> h*. Similarly, the initial condition for *c*_2_(*x, t*) is given by *c*_2_(*x*, 0) = *β >* 0 for |*x* − *x*_0_| *≤ h*, and *c*_2_(*x*, 0) = 0 for |*x* − *x*_0_| *> h*. This initial condition specifies that both subpopulations vanish outside an interval of width 2*h* centred at *x* = *x*_0_. Within this interval, the density of each subpopulation takes on a positive, constant value. For our approach, we require *k*_1_ *>* 0 and *k*_2_ *>* 0. The transformation and inverse of the transformation matrix are given by

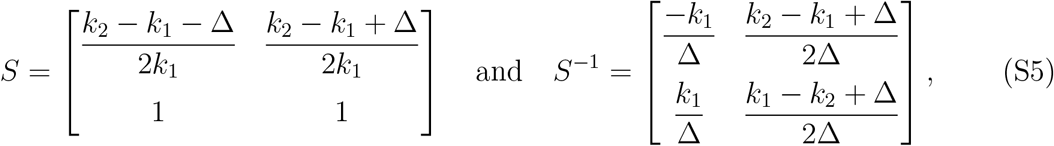

Where 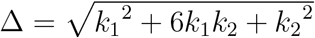. With these ingredients, we can write Equation (S4) in terms of the uncoupled variables *b*(*x, t*) = (*b*_1_(*x, t*), *b*_2_(*x, t*))^*⊤*^ as,

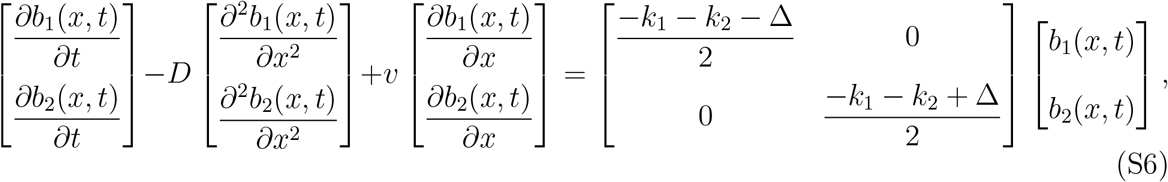

where we see that the square matrix associated with the source terms is now diagonal, which means that the two PDEs are uncoupled. Using *b*(*x*, 0) = *S*^−1^*c*(*x*, 0), the initial condition for *b*_1_(*x, t*) is given by *b*_1_(*x*, 0) = −*αk*_1_*/*Δ + *β*(*k*_2_ − *k*_1_ + Δ)*/*(2Δ) for |*x* − *x*_0_| *< h*, and *b*_1_(*x*, 0) = 0 for |*x* − *x*_0_| *> h*. Similarly, the initial condition for *b*_2_(*x, t*) is given by *b*_2_(*x*, 0) = *αk*_1_*/*Δ + *β*(*k*_1_ − *k*_2_ + Δ)*/*(2Δ) for |*x* − *x*_0_| *< h*, and *b*_2_(*x*, 0) = 0 for |*x* − *x*_0_| *> h*. With these initial conditions, each decoupled PDE can be solved using a standard integral transform approach [2] to give

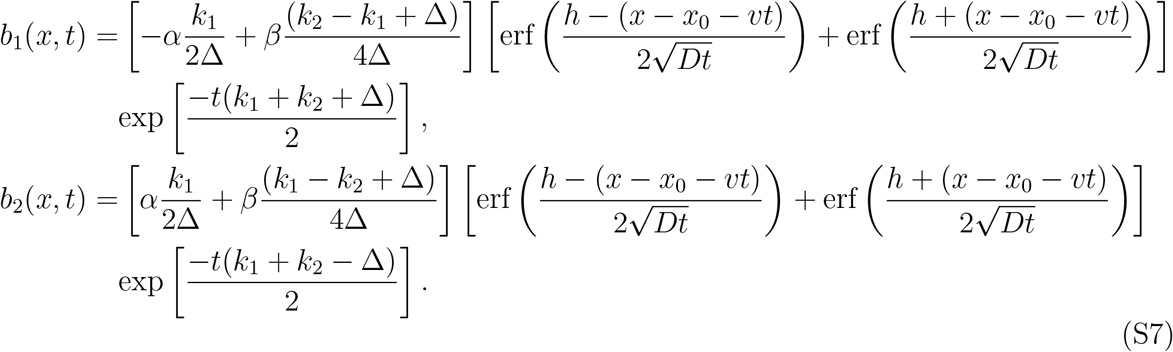

Using the relation *c*(*x, t*) = *Sb*(*x, t*) the exact solution to the coupled system of PDEs (Equations (2)–(3) in the main document) is

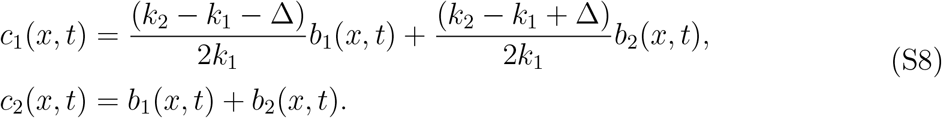

These solutions are strictly valid on the infinite domain, −*∞ < x < ∞*. In practice, we will apply these solutions on a finite domain, 0 *≤ x ≤ L*. In the usual way [3], choosing *x*_0_ and *L* to be sufficiently large that the density profiles for *c*_1_(*x, t*) and *c*_2_(*x, t*) remain close to zero near *x* = 0 and *x* = *L* allows us to use an exact infinite domain solution to approximate the solution of a closely-related finite domain problem [2].

## S2 Numerical solution of surrogate PDE for Case 2

Here we solve the continuum-limit PDE for the interacting random walk model (Equation (9) in the main document) using a numerical method-of-lines approach. To implement this approach we spatially discretise the PDE model using standard finite difference approximations for the spatial terms and then solve the resulting system of coupled ordinary differential equations (ODEs) in time using the DifferentialEquations.jl package in Julia to take advantage of automatic time stepping and temporal truncation error control [4].

We solve Equation (9) on 0 *< r < R*, with the initial condition given by *c*(*r*, 0) = *c*_0_ *>* 0 for *r < r*_0_ and *c*(*r*, 0) = 0 for *r > r*_0_. This initial condition specifies that the population vanishes outside a circle of radius *r*_0_ centred at the origin. Within this circle, the population density assumes a positive, constant value *c*_0_. We discretise the spatial terms on a uniform grid with grid spacing *δ >* 0, such that *c*(*r*_*n*_, *t*) = *c*_*n*_ for *n* = 1, 2, 3, …, *N*, where *r*_*n*_ = (*n* − 1)*δ*. At *r* = 0 and *r* = *R*, corresponding to mesh points *r*_1_ and *r*_*N*_, respectively, we impose zero-flux boundary conditions to give

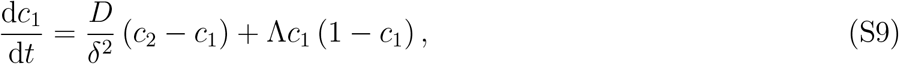

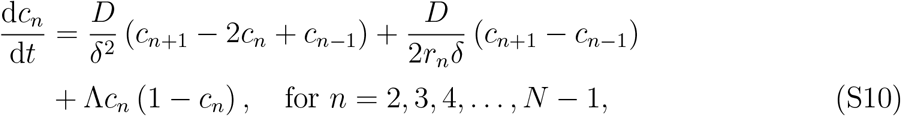

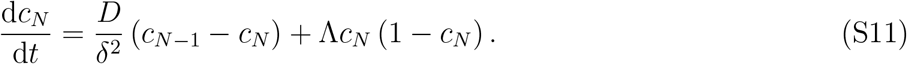

We solve this system of ODEs using Heun’s method within the DifferentialEquations.jl package in Julia [4]. All results presented in this work correspond to *δ* = 0.5. To ensure our results are grid independent, we considered a number of test cases and checked that numerical results with *δ* = 0.5 were indistinguishable from results with *δ* = 0.25.

## S3 Model validation

In this work, we employ two computationally efficient continuum-limit approximations of two types of RWMs. For the non-interacting RWM with two subpopulations, we use a system of PDEs (Equations (2)–(3) in the main document). For the interacting RWM with a single population, we use a two-dimensional (2D) generalisation of the well-known Fisher– Kolmogorov equation in radial coordinates (Equation (9) in the main document). Here, we present evidence demonstrating that the solutions of the surrogate PDE models give an accurate approximation of appropriately averaged data from the relevant RWM.

### S3.1 Case 1 validation

To validate the surrogate PDE for case 1 (Equation (2)–(3) in the main document), we compare exact solutions of the PDEs given by Equation (S8) with count data obtained from simulations of the noninteracting RWM. Since the solution of the PDE models gives us non-dimensional densities, *c*_1_(*x, t*) and *c*_2_(*x, t*), and the RWM simulations give count data, We compare the count data with the quantities *J × c*_1_(*x, t*) and *J × c*_2_(*x, t*) to give a direct comparison, noting that the PDE solution does not capture any fluctuations. To accurately compare noise-free PDE solutions with the noisy count data, we deliberately reduce stochastic variability in data from the non-interacting RWM by averaging the count data over 100 identically prepared realisations of the discrete model. Our comparison involves identical parameters, initial conditions, boundary conditions, and simulation time as in Case 1. In summary, the parameters of the simulation are (*α, β, h, x*_0_, *D, v, k*_1_, *k*_2_)^*⊤*^ = (0.5, 0.2, 19.5, 99.5, 0.25, 0.25, 0.02, 0.03)^*⊤*^ and the comparison is made at *t* = 100 on a large but truncated domain.

We present snapshots of a single realisation of the RWM simulation at the initial placement and after *k* = 100 time steps in Figure S1(a)–(b). The comparison in Figure S1(c)–(d) confirms that the same initial condition is used in both the PDE model and the RWM. The comparison after 100 time steps for both subpopulations is shown in Figure S1(e)–(f) which demonstrates that the solution of the surrogate PDE model accurately describes the output of the RWM when care is taken to minimise stochastic fluctuations.

**Figure S1.**
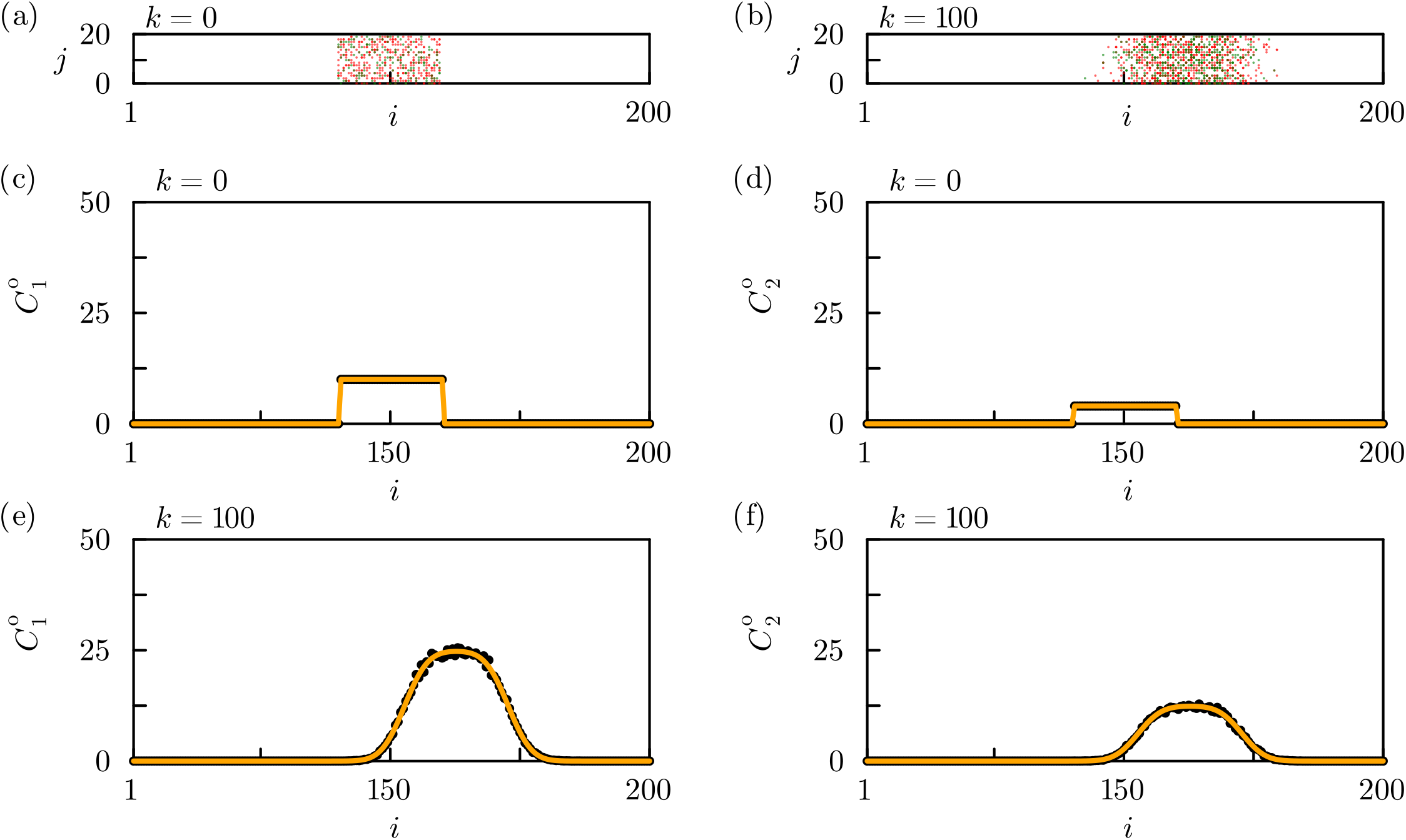
Validation of the surrogate PDE model for Case 1. he PDE solution is evaluated using (*α, β, h, x*_0_, *D, v, k*_1_, *k*_2_)^*⊤*^ = (0.5, 0.2, 19.5, 99.5, 0.25, 0.25, 0.02, 0.03)^*⊤*^. (a)–(b) Snapshots of the agent distribution. (c)–(f) Averaged count data from the non-interacting RWM over 100 realisations (black dots), superimposed with the corresponding PDE solution (solid orange line). (c) and (e) Comparison for subpopulation 1. (d) and (f) Comparison for subpopulation 2.

### S3.2 Case 2 validation

To validate the PDE for Case 2 (Equation (9) in the main document), we compare the solution of the PDE with count data obtained from a simulation using the interacting RWM. The PDE model is solved numerically as detailed in Section S2. The RWM is simulated using the same parameters, initial conditions, boundary conditions, and simulation time as in Case 2b of the main document. In summary we specify (*r*_0_, *D*, Λ, *c*_0_)^*⊤*^ = (20, 0.25, 0.01, 0.8)^*⊤*^ and compare the solutions at *t* = 600 for 0 *< r <* 100. Under these conditions we will now explain how we check that the solution of the surrogate PDE model provides a good prediction of the computationally expensive RWM.

We validate the PDE model by comparing the radial and column count data. For the radial count data, we consider the count data as a function of radius, *r*. The solution of the PDE, *c*(*r, t*), represents the population density at radius *r* and time *t*. In contrast, the RWM simulation can be summarised by the variable *C*^*\**^(*i, j, k*), which gives the number of agents at site (*i, j*) after *k* time steps in a single realisation. To convert the simulation result into a radial density profile, we superimpose *Q* equally-spaced concentric circles on the lattice. Each circle, shown in Figure S2(b) with *Q* = 20, is centred at (*i, j*) = (100, 100), and the *q*th circle has radius *r*_*q*_ = 100*q/Q* for *q* = 1, 2, 3, …, *Q*. For *q ≥* 2, the density at radius *r* is approximated by counting the number of agents within the annulus *r*_*q*−1_ *< r < r*_*q*_. For the smallest concentric circle, *q* = 1, the density is approximated by counting agents in the circular region *r < r*_1_. This process yields *Q* equally-spaced estimates of the agent counts as a function of *r*. To reduce the impact of stochastic fluctuations, we average these counts over 100 identically prepared realisations of the discrete model, noting that averaged count data can lead to non-integer values. We compare this averaged count data with the solution of the surrogate PDE (Equation (9) in the main document) in terms of the quantity (2*πr*Δ) *× c*(*r, t*), where *c*(*r, t*) is the solution of the surrogate PDE model. The factor 2*πr*Δ is the area of the annulus of thickness Δ, centred at radial position *r*. In our simulations, we have Δ = 100*/Q*.

For the column count data, we consider the count data as a function of the horizontal position, *x*. The count data are obtained from the RWM simulation, consistent with the approach described in Section 2.3 of the main document, except that the data are averaged over 100 identically prepared realisations. To ensure consistency with the units of the RWM simulation, the PDE solution is rescaled by *H × c*(*r, t*), which represents the agent count along the radius. As in Section 2.3, we take *H* = 20. By symmetry, we have *r* = |*x* − *x*_0_|, where *x*_0_ denotes the *x*-coordinate of the centre of the spreading population. The scaled PDE solution can therefore be expressed as a function of horizontal position, *C*°(*x, t*).

Our discrete-continuum comparison is performed for the radial count data, *C*°(*r, t*), and column count data, *C*°(*x, t*), summarised in Figure S2. The initial distribution of agents is given in Figure S2(a), and the resulting distribution of agents after *k* = 600 time steps is given in Figure S2(b) where we see the impact of combined migration and proliferation as the population spreads away from the initial region containing the agents. Note that the impact of proliferation is clear as the number of agents on the lattice after *k* = 600 time steps has clearly increased, and the density of agents towards the centre of the population, near (*i, j*) = (100, 100) where *r* = 0, is high. In this region the population forms an almost confluent monolayer, which is close to the maximum packing density in the interacting RWM. The distribution of radial count data, *C*°(*r, t*), as a function of radial position obtained from the discrete model is superimposed on count estimates using the surrogate PDE model plotted in Figure S2(c)–(d) for *t* = 0 and *t* = 600, respectively. The distribution of column count data, *C*°(*x, t*), as a function of horizontal position obtained from the discrete model is superimposed on count estimates using the surrogate PDE model plotted in Figure S2(e)–(f) for *t* = 0 and *t* = 600, respectively. Comparing the discrete count data with the distribution of count data from the surrogate PDE solution confirms that the solution of the surrogate PDE provides a good approximation of RWM in terms of agent counts where care has been taken to construct stochastic count data with minimal fluctuations.

**Figure S2.**
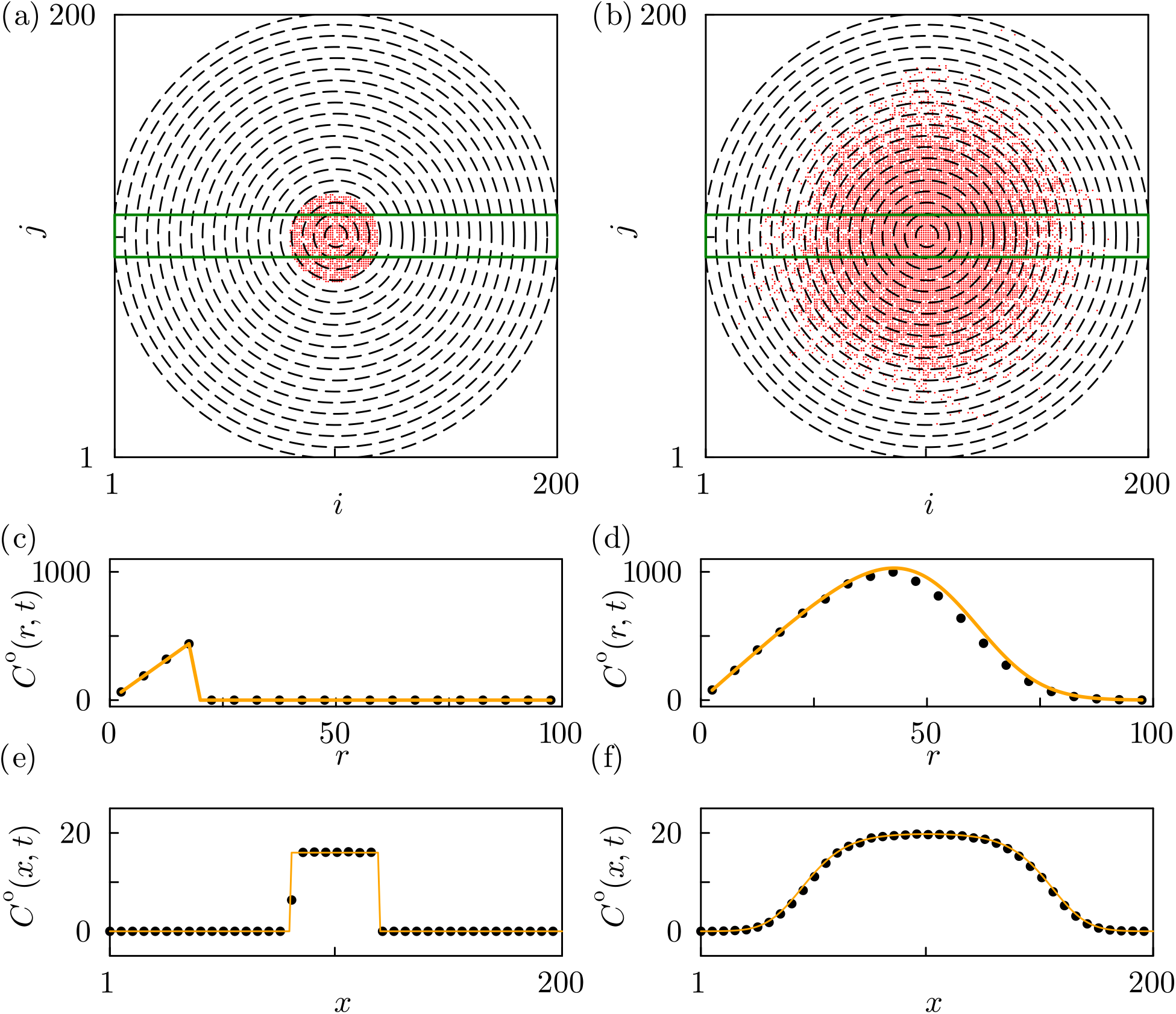
Validation of the surrogate PDE model for Case 2. The PDE is solved using (*r*_0_, *D*, Λ, *c*_0_)^*⊤*^ = (20, 0.25, 0.01, 0.8)^*⊤*^. (a)–(b) Snapshots of the agent distribution at *k* = 0 and *k* = 600, respectively. Each snapshot is superimposed with 20 concentric circles (black dashed lines) and a rectangular box (green line), which are used to construct the radial and column count data from the stochastic model, respectively. (c)–(f) Discrete simulation results (black dots) superimposed with the solution of the surrogate PDE model (solid orange line). (c)–(d) Radial count data comparisons at *t* = 0 and *t* = 600, respectively. (e)–(f) Column count data comparisons at *t* = 0 and *t* = 600, respectively.

## S4 Additional Results: Case 1 over a longer timescale

Results for estimation, identifiability and sensitivity analysis for Case 1 in the main document are based on data captured after a relatively short simulation, *k* = 100. Here we repeat the exact same analysis for data collected after a longer duration of time, *k* = 200. The data for this additional case is summarised in Figure S3 where these simulations are precisely the same as used in the main document except that data are collected after *k* = 200 time steps instead of *k* = 100 time steps.

New results in terms of parameter estimation, parameter identifiability and sensitivity analysis via profile-wise predictions are summarised in Figure S4 for the new data collected after *k* = 200 time steps. Comparing these additional results with those in the main document (Figure 4) for the shorter time scale simulation reveals that there is no striking difference in terms of parameter estimation, parameter identifiability and sensitivity analysis in this case.

**Figure S3.**
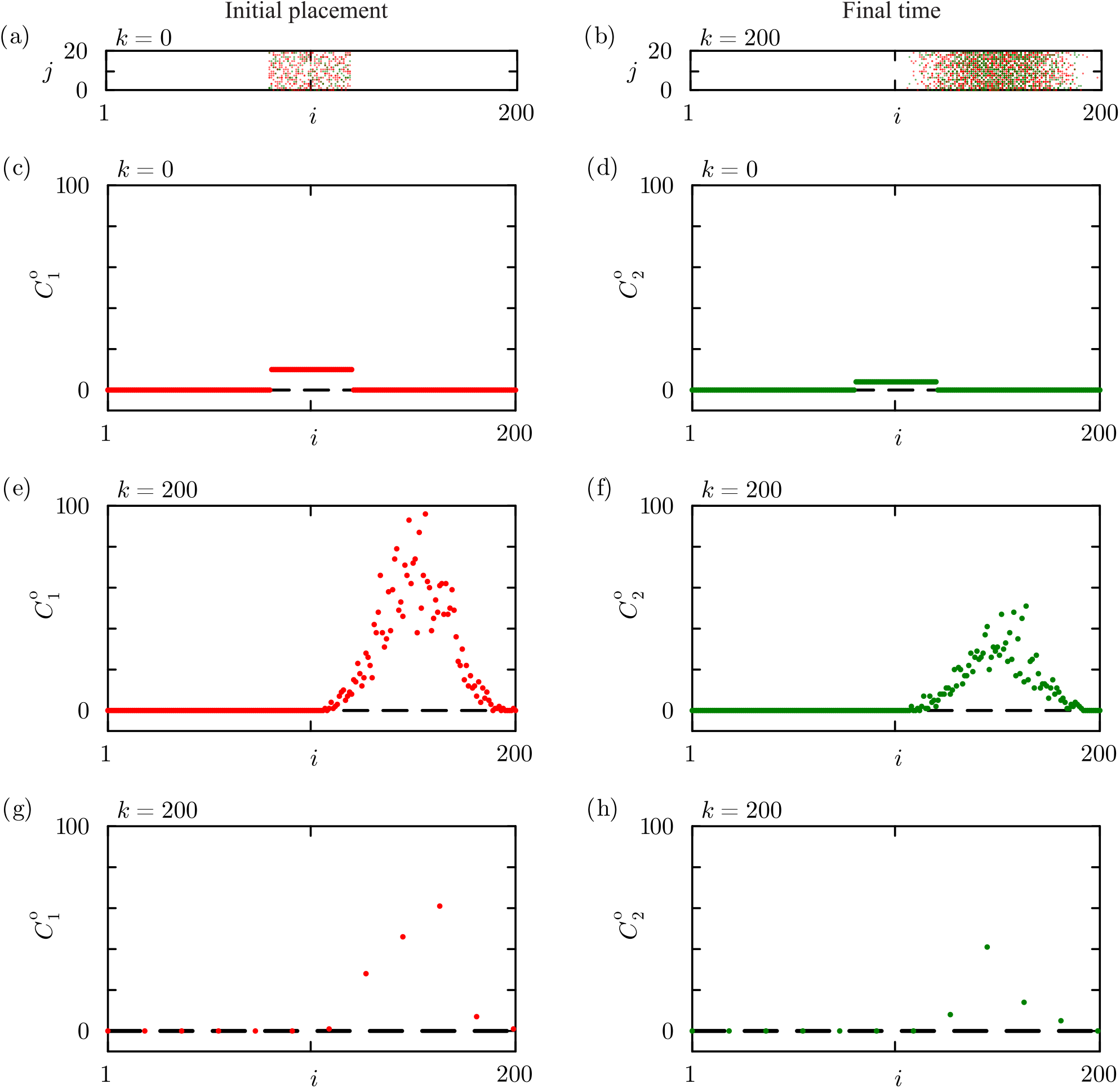
Snapshots of agent distributions and count data from the non-interacting RWM with (*P, ρ, λ*_1_, *λ*_2_)^*⊤*^ = (1, 0.5, 0.02, 0.03)^*⊤*^. (a)–(b) Snapshots of agent distribution. (c)–(f) Full count data. (g)–(h) Subsampled count data obtained from data in (e)–(f), respectively. Horizontal dashed lines in (e)–(h) show the 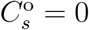 lower bound for the count data. Results correspond to *I* = 200 and *J* = 20.

**Figure S4.**
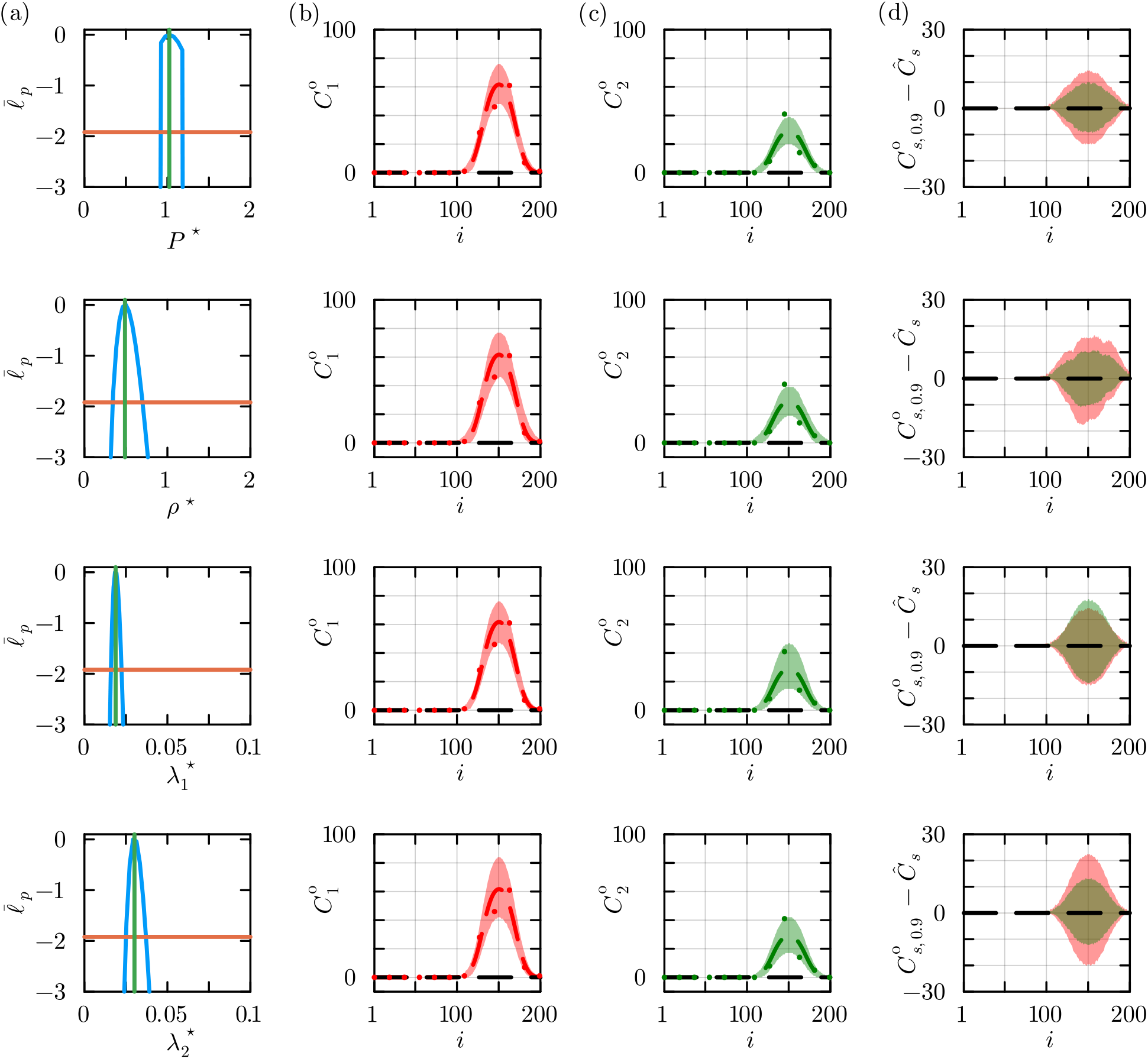
Estimation, identifiability and sensitivity results for the non-interacting RWM. (a) Univariate profiles for 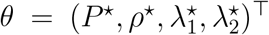 indicate the MLE (vertical green line) and the 95% asymptotic threshold (horizontal orange line). The MLE is 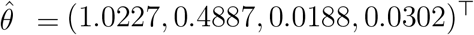. (b)–(c) Profile-wise prediction intervals for *C*_1_ (shaded red) and *C*_2_ (shaded green) for each parameter superimposed with count data (green dots), and the solution of the PDE evaluated at the MLE for *C*_1_ (dashed red) and *C*_2_ (dashed green). (d) Difference between confidence set and the solution of the mathematical model evaluated at the MLE for *C*_1_ (shaded red) and *C*_2_ (shaded green).

